# Orthographic training finetunes a neuronal letter position code in primate visual cortex

**DOI:** 10.64898/2025.12.01.691513

**Authors:** Aakash Agrawal, Shubhankar Saha, Surbhi Munda, SP Arun, Stanislas Dehaene

## Abstract

Reading is an acquired skill that enhances brain activity in human visual cortex and is thought to repurpose preexisting ventral visual circuitry for the fast parallel processing of letter strings. However, the neuronal code underlying position-invariant written word recognition remains elusive. Here, we examined this issue in a macaque monkey model before and after orthographic training. Based on prior simulations of reading acquisition in a convolutional neural network model of the ventral visual pathway, we hypothesized the existence of neurons sensitive to both specific letters and their ordinal position within the word, which should be enhanced by orthographic training. By wirelessly recorded neural activity from the inferior temporal (IT) cortex of macaque monkeys trained on orthographic tasks over five successive days, we indeed discovered IT neurons tuned to letters and sensitive to either ordinal or retinotopic positions. Ordinal units existed prior to training, but their responses were enhanced after training, matching with behavioral improvements in word recognition. These findings support the neuronal recycling hypothesis and demonstrate, at the single-cell level, how reading refines pre-existing neural circuits to facilitate fluent word recognition.

## Introduction

Reading is a uniquely human cognitive skill that transforms arbitrary visual shapes into meaningful linguistic constructs, facilitating communication and the transmission of knowledge across generations. Unlike face recognition or spoken language comprehension, reading is a cultural invention and an optional skill whose acquisition, following extensive training and practice, leads to significant neuroplastic changes in the brain (Dehaene et al., 2015). One of the most profound changes associated with literacy acquisition is the emergence of the Visual Word Form Area (VWFA) in the left ventral occipitotemporal cortex, a visual region selective for letter strings in the learned orthographic system (Baker et al., 2007; Cohen et al., 2002; Dehaene-Lambertz et al., 2018). This region acquires its selectivity only in literate individuals – otherwise, in pre-reading children as well as illiterate adults, it participates in the visual recognition of visual objects and faces (Dehaene, Pegado, et al., 2010; Pegado et al., 2014). This finding exemplifies the brain’s capacity for *neuronal recycling*, where pre-existing neural circuits are partially repurposed for new functions (Dehaene, Pegado, et al., 2010; Dehaene & Cohen, 2007; Gomez et al., 2019; McCandliss et al., 2003).

The neuronal recycling hypothesis posits that the VWFA emerges by repurposing a small part of the ventral visual cortex originally dedicated to object and shape recognition. This repurposing leverages this pathway’s inherent capacity to process complex visual stimuli invariantly across changes in size, position, font, and other visual transformations (Bao et al., 2020; DiCarlo & Cox, 2007; Freiwald & Tsao, 2010; Grill-Spector & Malach, 2004; Kourtzi & Kanwisher, 2001; Vighneshvel & Arun, 2015). Indeed, in literate adults, the VWFA exhibits a degree of invariance to variations in visual presentation, allowing for consistent recognition of written words despite superficial differences (Dehaene et al., 2001, 2004). Beyond its visual properties, the VWFA is also known to encode the statistical properties of words such as letter or bigram frequency, and its response gradually decreases as the similarity of the target letter string to existing words decreases (Binder et al., 2006; Vinckier et al., 2007; Woolnough et al., 2021; Zhan et al., 2023). Since they depend on the target language, those properties must be acquired, again capitalizing on the capacity of ventral visual cortex to get attuned to the statistics of its inputs, which has been documented for other visual stimuli in both humans and monkeys (Li & Dicarlo, 2012; Logothetis et al., 1995; Miyashita, 1988).

Investigating the neural basis of orthographic processing at the neuronal level in humans is challenging due to limitations of non-invasive imaging techniques. Non-human primates offer a valuable model for studying these mechanisms, as they share similar object recognition behavior and neural representations with humans, particularly within the inferior temporal (IT) cortex, which is involved in high-level visual recognition (DiCarlo et al., 2012; Tanaka, 1996) and homologous to the human ventral visual pathway (Bao et al., 2020; Tsao et al., 2008). Although the semantic and phonological components of reading are beyond their reach, monkeys can perform orthographic tasks that do not require linguistic comprehension and learn to discriminate letter strings based on their statistical regularities. For instance, Grainger et al. (2012) demonstrated that baboons could learn to distinguish English words from non-words by recognizing frequent letter combinations, with behavioral signatures similar to orthographic processing in human (Ziegler et al., 2013). Moreover, when untrained monkeys were exposed to letter strings, the activity of neurons in the IT cortex could predict the error patterns of baboons with the same stimuli (Rajalingham et al., 2015). The neuronal responses of untrained monkeys to letter strings can even be used to solve CAPTCHA challenges, suggesting an invariance to visual transformations similar to humans (Katti & Arun, 2022). Consistent with IT responses to multiple objects (Sripati & Olson, 2010; Zoccolan et al., 2005), the firing of IT neurons to letter strings is well predicted by the sum of its responses to individual letters (Katti & Arun, 2022). These findings suggest that the primate visual system possesses a pre-existing architecture capable of supporting orthographic processing and which can be enhanced through training—a principle consistent with neuronal recycling.

Despite these insights, the effects of orthographic training on individual IT neuron responses remain largely unexplored. In humans, the impact of training in reading has been explored solely through comparisons of literate versus illiterate subjects, using non-invasive brain-imaging techniques such as fMRI or M/EEG which are unable to resolve single-neuron codes (Brem et al., 2010; Dehaene, Pegado, et al., 2010; Dehaene et al., 2015; Dehaene-Lambertz et al., 2018; Feng et al., 2020; Maurer et al., 2006; Pegado et al., 2014). In monkeys, while we know of no training studies with letter strings, previous studies have shown that familiarity with visual stimuli can lead to a decrease in overall neural activity and enhanced selectivity for specific object features or whole objects in the IT cortex (Baker et al., 2002; Logothetis et al., 1995; Meyer et al., 2014). This raises the question of whether similar neural plasticity occurs during the acquisition of orthographic skills, particularly concerning the encoding of letter identities and their ordinal positions. Here, we investigate this issue by training monkeys to recognize letter strings across a short delay, and examining neural responses before and after training.

Our research is particularly targeted at the issue of how letters and their order are encoded. Selectivity to letter order is a critical aspect of fluent reading since the same 26 letters are used to make up thousands of different words. Accurate representation of letter position is essential for distinguishing words such as “FROM” and “FORM”, which contain identical letters in different order. To explain how the brain encodes letter sequences, cognitive models of visual word recognition have proposed several coding schemes such as position-specific letter detectors, open-bigram representations, and spatial coding models (Grainger & Van Heuven, 2004; McCloskey et al., 2013; Norris, 2013; Whitney, 2001). Despite their theoretical appeal, these models remain untested against neural data.

In the past twenty years, one prominent model proposed that words are encoded by a list of their letter bigrams—ordered pairs of letters such as “E left of R”, possibly with intervening letters (Dehaene et al., 2005; Grainger & Van Heuven, 2004; Vinckier et al., 2011). However, recent empirical work suggested an alternate view where the visual representation space of words can be modelled as a linear combination of its individual letters at specific positions, without the need to evoke bigrams (Agrawal et al., 2019, 2020). Similar findings were conclusions were obtained in recent simulations of deep neural network models: after training in word recognition, word-selective units emerged in the deeper layers, whose response could largely be predicted using a letter by position model (Agrawal & Dehaene, 2024; Hannagan et al., 2021). Specifically, we “recycled” a previously proposed convolutional neuronal model of visual recognition by training it to recognize one thousand different written words in various fonts, cases and sizes. After training, we discovered that some of the units had become specialized for alphabetic stimuli, including specific letters and ordinal positions. These units, in the early layers, behaved as “space bigrams“: they jointly encoded the presence of a letter and of a blank space at a specific approximate distance to its left or to its right. For instance, units could respond to “A immediately right of space” (i.e. A in first ordinal position), or “E at some distance left of space” (i.e. E at a position near the end of the word, but not last). We found that, across the network hierarchy, these units were progressively pooled to form representations that encoded solely the ordinal positions of letters within words, regardless of their actual retinotopic position. This model thus could explain how a neural network achieves positional invariance, and why the middle letters can be trasnposed or partially srcamlbed with only a limited impact on word recognition (Perea & Lupker, 2003; Schoonbaert & Grainger, 2004; Ziegler et al., 2013). Recently, we used high-field 7T fMRI and MEG to obtain indirect evidence for such ordinal representations in the VWFA and along the anterior regions of the human ventral visual pathway of literate humans (Agrawal & Dehaene, 2025). However, at the single-unit level, the model is highly speculative, and empirical evidence for the existence of letter- and ordinal-position-selective neurons in the visual system is currently lacking.

To address these questions, we recorded wirelessly from IT neurons before, during, and after two macaque monkeys were trained on an orthographic delayed match-to-sample (DMS) task over five successive days. This training period may seem short, but note that the monkeys were very familiar with the DMS task, as they had been trained extensively to perform it with picture stimuli – we merely transferred them to letter string stimuli (Jacob et al., 2021). Furthermore, the target stimuli for the DMS task were only 36 written “words” made only of 6 letters (3 consonants and 3 vowels), thus modeling how readers in a certain script are trained to focus on specific letter shapes and spellings. Before and after such training, we measured neuronal activity to a greater variety of letter combinations using a passive fixation task with rapid serial visual presentation (RSVP), allowing us to assess changes in neural coding independent of task performance. In this study, we focus on the data from the RSVP task recorded before and after training, which has higher SNR due to repeated stimulus presentation and allows to address the changes in neuronal responses with training.

To anticipate on the results, we found that, as in previous studies,, even before training, many IT neurons were tuned to letters at contralateral locations (Katti & Arun, 2022; Rajalingham et al., 2020). Using stimuli carefully designed to probe positional coding, we further identified a subset of neurons that selectively encoded the ordinal position of letters within words. With training, these representations evolved to reflect behavioral performance. Behavioral model fits revealed that Monkey 1 processed letter strings in a position-invariant manner and, accordingly, exhibited enhanced ordinal position coding in IT neurons after training. Monkey 2, however, did not improve in the delayed match-to-sample task and showed enhanced retinotopic coding.

Complementary simulations using convolutional neural network (CNN) models of reading revealed a similar pattern. In the response to the same stimuli as the monkeys, ordinal coding units were found in deeper layers of the CNNs, and were enhanced in literate (trained) compared to illiterate (untrained) networks, mirroring the neural organization observed in the monkeys’ IT cortex. Together, these findings suggest that hierarchical processing in both artificial and biological systems gives rise to a compositional letter X ordinal position code essential for orthographic processing (Agrawal & Dehaene, 2024; Hannagan et al., 2021; Yamins & DiCarlo, 2016; Yin et al., 2023).

## Results

We trained two macaque monkeys on an orthographic delayed match-to-sample task (delayed match to sample) while wirelessly recording the activity of neurons in their inferior temporal (IT) cortex. The stimulus set consisted of four-letter strings, most of which were composed of six visually distinct letters: A, E, O, P, L, and W (3 consonants and 3 vowels). These letters were combined to generate 36 stimuli designated as “words” and following two specific consonant-vowel (C-V) structures, either CVCV or VCVC (e.g. WOLE, ALAP). Our choice of words ensured uniform occurrence of each letter across all four positions, as well as balanced bigram pairings within the words (see full list in Methods). To investigate position invariance and distinguish putative neurons with retinotopic versus word-centered receptive fields, these words were presented at five different positions relative to fixation: -2, -1, 0, +1, and +2 letters from the center (Figure 1C). In addition to those words, we also presented nonword stimuli designed to vary in their similarity to the words and to probe different aspects of orthographic processing. There were 6 types of nonwords that ranged from strings composed of entirely unfamiliar letters to those involving letter transpositions and substitutions (Figure ID; see Methods for details)

**Figure 1:**
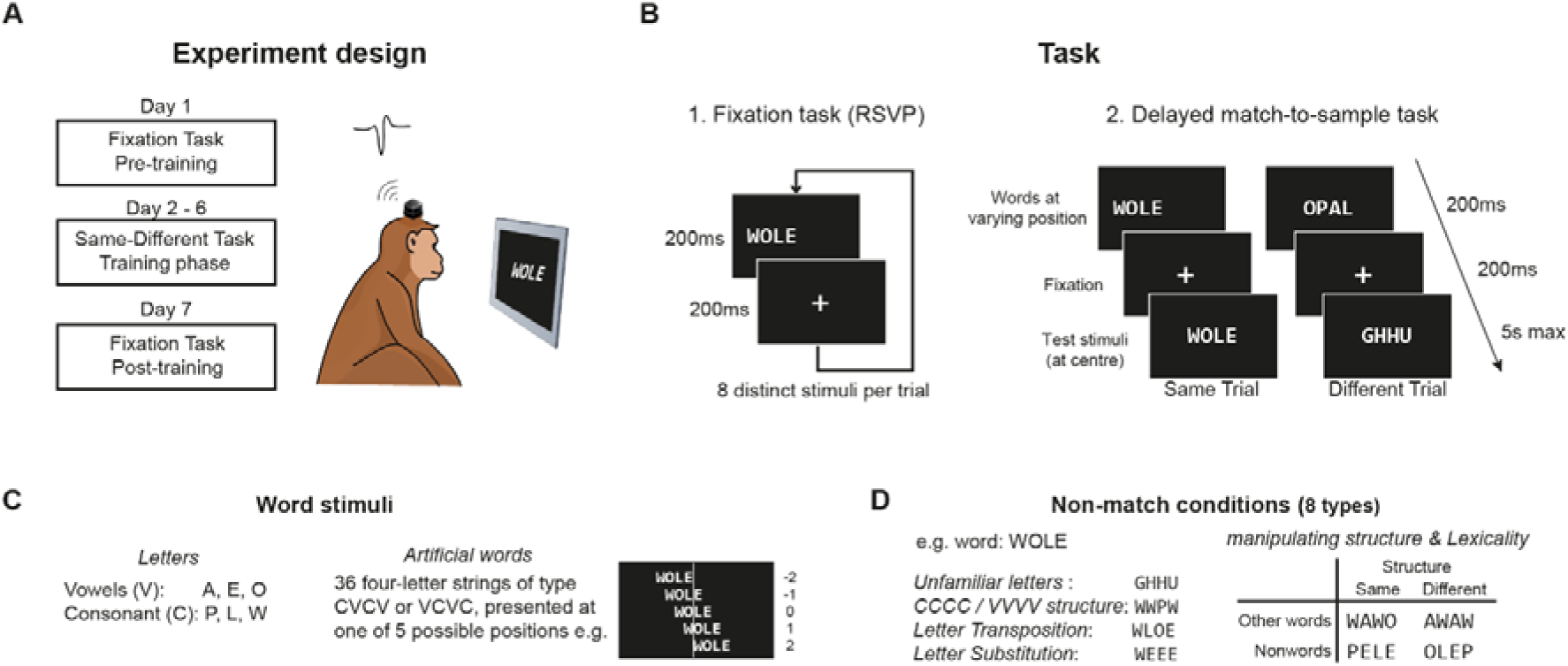
Design for fixation task and delayed match-to-sample task. (A) Pre-training, training and post-training phases of the study. Monkeys performed RSVP fixation before and after training session, which lasted for 5 days while the neural data were recorded wirelessly. (B) In the fixation task, stimuli were presented while monkeys maintained their fixation without making active responses. In the delayed match-to-sample task, a sample stimulus was displayed for 2001ms, followed by a 2001ms blank screen, then a test stimulus appeared until response. Monkeys indicated whether the test stimulus matched the sample. Sample words were presented at five positions relative to fixation: -2, -1, 0, +1, and +2 letters from the center, testing position invariance and retinotopic encoding. (C) Example word stimuli used in this study. They were always centered vertically and shifted along the horizontal axis. (D) Example non-match stimuli used in this study. For each word, there were 8 types of non-match stimuli, including 6 non-word categories, that varied in structure, lexicality and visual similarity to words.

During training, the monkeys performed a delayed match-to-sample (DMS) task (Figure 1B). First, one of the 36 words was presented as sample for 200 ms at one of the five positions relative to fixation, followed by a 200 ms blank screen, and then a test stimulus, which could be a word or nonword, appeared and remained at the center until the monkey made a response. The monkeys were required to indicate whether the test stimulus matched the sample word or not. On half the trials, the test and the sample matched (although they could occupy different retinotopic positions, to promote the emergence of ordinal coding). On the other half of the trials, a variety of mismatches were presented: from easy cases composed of entirely different letters (analogous to the “false-font” condition in humans) to more challenging cases in which the test stimulus was another word/nonword with the same or a different structure, and the most difficult cases involving middle-letter transpositions or substitutions. Training sessions were conducted over five consecutive days, with new non-words randomly generated each day to avoid any memorization. To assess neural responses independently of task demands, the monkeys also performed a RSVP fixation task on pre-and post-training days (Figure 1B), where stimuli were presented briefly for 200ms with a 200 ms fixation interval (SOA=400 ms) while monkeys had to maintain fixation for 8 consecutive trials without making any saccades to receive juice reward.

### Behavioral evidence of learning

We observed above-chance performance in the delayed match-to-sample task for both monkeys across all five days of training, indicating that monkeys indeed learnt the task. In monkey 1, performance improved steadily over time. The d-prime (d1) steadily increased from 0.30 on day 1 to 0.79 on day 5, reaching its peak by day 4 (Figure 2A). Response times (RTs) for both match and non-match conditions decreased with training suggesting enhanced processing speed (Figure 2B). The enhanced performance was statistically significant (p < 0.0005) as estimated by assessing the proportion of times day 1 was higher than day 5 across 100 bootstrap. Conversely, monkey 2 did not exhibit significant changes in performance across the training days; Unlike monkey 1, its best performance was observed on the first day (d’ = 0.48), with d1 values remaining relatively stable thereafter (average d1 = 0.38).

**Figure 2:**
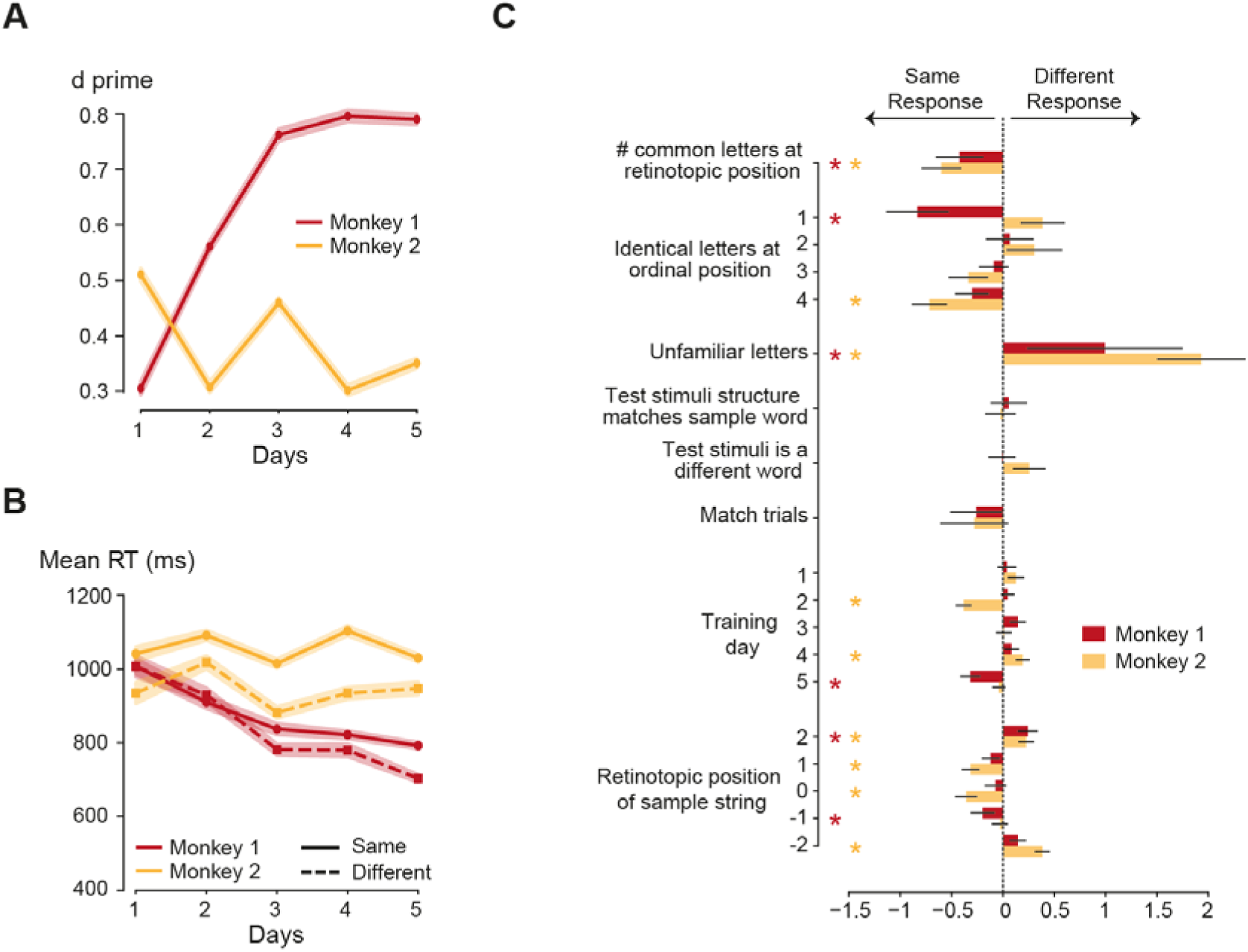
Behavioral Performance. (A) Performance in discriminating match and non-match trials (d1 values) over five days of training for both monkeys. Shaded error bars represent one standard error of mean across 100 bootstrap (sampling with replacement) estimates. (B) Response times for correct match (solid) and non-match (dashed) conditions across training days for monkey 1 (red) and monkey 2 (orange). Shaded error bar indicates standard deviation across 100 bootstrap estimates. (C) Logistic regression coefficients illustrating factors influencing “same” and “different” responses for monkey 1 (red) and monkey 2 (orange). Error bars indicate standard deviation across 100 bootstrap estimates. Asterisks indicate statistical significance after Benjamini-Hochberg FDR correction (alpha < 0.05). P-values were estimated from the bootstrap distribution (n = 1000).

On average, RTs were faster for non-match conditions compared to match conditions in both monkeys (Figure 2B). This trend may be expected given that a non-match can be detected based on a mismatch in any of the letters. Overall task accuracies for monkey 1 increased from 56% correct on day 1 to 65% on day 5. For monkey 2, accuracies remained relatively constant, ranging from 60% on day 1 to 57% on day 5 (see Figure S1 for accuracy values separately for match and non-match conditions). Both monkeys exhibited the lowest error rates when the sample stimulus was presented at the center of fixation. Error rates increased as the sample position moved further from the center, indicating decreased performance for peripheral stimuli (Figure S2A), similar to human readers (Pelli & Tillman, 2008). No significant differences in RTs were observed across sample positions (Figure S2B).

Across the different non-match conditions, both monkeys performed best (i.e., lower error rates and faster RTs) in the unfamiliar letter condition, where all letters in the test stimulus were unusual and necessarily different from the sample. For monkey 1, error rates were highest for nonwords that were very close to words, involving only transpositions and substitutions of internal letters. Other nonword conditions showed intermediate performance levels. In monkey 2, except for the easy rejection of unfamiliar letters, performance was relatively uniform across other nonword types (Figure S3). Note that both monkeys did not make more errors for internal transpositions compared to letter substitutions, i.e. they did not exhibit a letter transposition effect, unlike human readers (Perea & Lupker, 2003; Schoonbaert & Grainger, 2004; Ziegler et al., 2013). This could be due to insufficient training, as there is evidence that the effect depends on reading expertise (for discussion, see Hasenäcker & Schroeder, 2022; Ziegler et al., 2014).

To further quantify the monkeys’ decision strategies, we trained a logistic regression model to predict their responses (i.e., same or different) across all five days of training (see Methods). The model achieved single-trial classification accuracies of 671±11% for monkey 1 and 621±11% for monkey 2, significantly above chance levels (50%). Visualization of the model coefficients revealed several interesting patterns (Figure 2C) on the strategy used by both monkeys to perform the delayed match-to-sample task. In both monkeys, as noted earlier, “different” responses were more likely when the test stimuli comprised letters that never appeared in the sample (unfamiliar letter condition). Conversely, “same” responses were more likely 1) when the sample stimulus was presented closer to the center of fixation; 2) at the retinotopic level, when there was a higher fraction of common letters at the same retinotopic position between the sample and test stimuli. 3) at the ordinal level, when the sample and test stimuli ended with the same letter (both monkeys) or started with the same letter (monkey 1). It is interesting that monkey 1, who progressed during the 5 training days, made errors when the first and last letters at ordinal position were the same, suggesting that he began to understand the need to encode the ordinal position of letters within the word, independently of the position of the whole word (and did not care about the two internal letters, which explains the lack of a transposition effect). Monkey 2, on the other hand, who did not progress, seem to focus only on the word-ending letter.

A similar pattern emerged when using regression models to predict response times for nonword conditions (Figure S4). The monkeys were faster to detect non-match conditions when all letters were different, and slower when there was a larger overlap in the number of common letters between the sample and test stimuli, particularly in conditions involving an identical initial letter for monkey 1 and an identical last letter for monkey 2.

### Average neural population responses in the fixation task

We wirelessly recorded from 182 (and 179) neurons in Monkey 1 during pre (and post) training respectively. Similarly, in monkey 2, we recorded from 194 (and 200) neurons during pre (and post) training. The electrodes were implanted in the inferior temporal cortex of both monkeys performing a fixation task (see Methods). Spike sorting was manually administered to estimate single unit activity wherever possible. For each neuron, firing rates were calculated using non-overlapping 20 ms time bins across all repetitions of a stimulus during which the monkey maintained its fixation. These firing rates were normalized using pre-stimulus activity (see Methods). For all further analysis we selected units with reliable activity as defined using a significant split-half correlation (see Methods).

On average, neural activity peaked around 60 ms post-stimulus when at least one letter was within the contralateral visual field (Figure 3A). When stimuli were presented entirely in the ipsilateral visual field, firing rate was much lower during the initial time window and only peaked later (∼250 ms). This delayed response is presumably due to corrective eye movements that brought part of the first letter into the contralateral receptive field (Figure S5). Eye-tracking data suggests that such saccades occurred when stimuli were presented on the extreme periphery, but that these eye movements averaged no more than about 0.5 to 1 letter width, potentially only bringing peripheral letters just slightly closer to the fovea. To preclude saccadic influences, subsequent analyses were limited to the first 250ms. Overall, neural responses to words varied gradually depending on the number of letters presented within the contralateral receptive field; the more letters that appeared in the contralateral field, the stronger the observed neural response (Figure 3A). Across different nonword conditions, neural responses were not significantly different from each other, except for the condition where all letters were unfamiliar and which led to higher firing (Figure 3A). This finding suggested that the neuronal population quickly adapted to frequent letters, which were indeed more frequent in the RSVP block even prior to training, and had higher responses to the rare unfamiliar letters.

**Figure 3:**
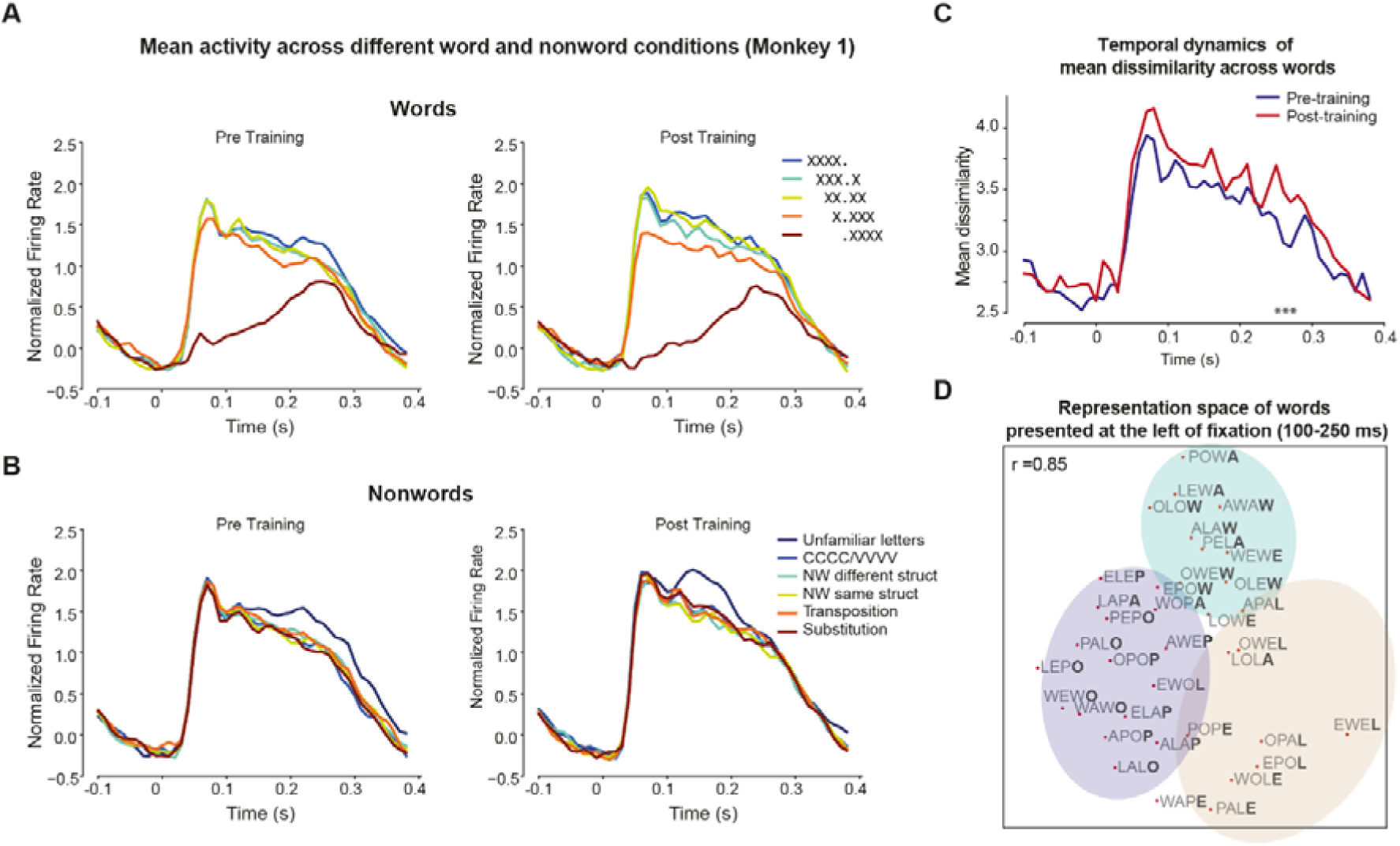
Average neural population responses and representational similarity analysis. (A) Average normalized neural response over time for words presented across different spatial position during pre (left and post (right) training. The neural responses were highest for the stimuli presented as the center, and minimal for stimuli that fell fully in the contralateral (right) hemifield. (B) Same as (A) but for different categories of nonwords. We observed a sustained increase in activity for unfamiliar letters post training compared to the other conditions. (C) Mean pairwise dissimilarity for word stimuli in Pre (blue) vs. post-training (red) sessions. Post-training responses show increased dissimilarity, particularly in the late time window (∼250 ms). Asterisks denote significant differences in mean dissimilarity (p < 0.05), assessed via a permutation test (N=1000 iterations). The null distribution was generated by shuffling neuron Firing Rates between Pre and Post sessions and recomputing the difference in mean dissimilarity at each time point. (D) Multidimensional scaling (MDS) plot of the neural representational space for words (pretraining 100-250ms) presented at the extreme left position, illustrating clustering based on the last letter and letter shape – letters “E” and “L” are clustered together; similarly, for “A” & “W” and “O” & “P”. The last letter of each word is highlighted to emphasize this clustering.

With training, neural activity increased for both words and nonwords (Figure 3B). Notably, the increase in activity for the Unfamiliar letters condition shifted to an earlier time and remained significantly different only during a brief period (100–200 ms). This shift could underlie the faster response times observed post-training for detecting non-matching stimuli. The response profiles in monkey 2 differed characteristically from those in monkey 1, yet we observed qualitatively similar patterns (Figure S6). In monkey 2, prior to training, there was no significant difference between the unfamiliar letters condition and other nonword conditions, although unfamiliar letters could still be reliably decoded from word responses during the late time window (Figure S14). Post-training, the neural response to the unfamiliar letters condition became more pronounced, mirroring the pattern seen in monkey 1 (Figure S6).

### Representational similarity analysis

Next, at each 201ms time step, we computed the representational dissimilarity matrix (RDM) across all 36 word stimuli using only the units with significant split-half consistency either during pre or post training. In monkey 1, the dissimilarity values increased post-training (Figure 3C)i, suggesting an overall scaling of the representational space and enhancing discriminability among different strings, as observed in literate humans (Agrawal et al., 2019). This increase in dissimilarity was much lower for monkey 2 (Figure S6) suggesting that improvements in orthographic task performance are closely linked to an increase in the scale and separability of neural representations in IT cortex.

We visualized the representational space for words presented at the extreme left position, as this contralateral condition elicited the highest neural activity. The RDM estimated using the 100-250 ms of neural data was embedded using non-metric multidimensional scaling (MDS). The resulting clustering was not based on the overall word structure (e.g., CVCV vs. VCVC). Rather, stimuli with the same last letter (closest to fixation) often grouped together, indicating a primarily retinotopic representation based on letters closer to fixation (Figure 3D). Additionally, letters with similar shapes (e.g., ‘O’ and ‘P’, ‘E’ and ‘L’, ‘A’ and ‘W’) were positioned closer to each other in the embedded space, suggesting shape-based encoding. Similar results were observed when considering all stimuli together (Figure S7), indicating that the observed patterns were consistent across different spatial configurations.

To further investigate the factors contributing to the neural representational space, we modelled it as a linear combination of pairwise dissimilarities based on word positions, retinotopic letter positions, and ordinal letter positions (Figure S8-S10). As expected, the first two factors—word position and retinotopic letter position—explained most of the variance in the data (Figure S9-S10). From the estimated model coefficients, we observed significant contributions from letters presented in the contralateral visual field. Additionally, there was a significant contribution from the first ordinal position in monkey 1 (Figure S9) and from the last ordinal position in monkey 2 (Figure S10), suggesting a slight but significant presence of ordinal position encoding.

### Single unit responses

While the population-level neural dynamics were primarily driven by retinotopic properties of the stimuli, we investigated whether individual neurons exhibited similar trends. Although most neurons encoded retinotopic positions and selectively responded to specific letter, there were also neurons that encoded ordinal letter positions even before training (Figure 4A), and most neurons exhibited mixed selectivity, responding to combinations of retinotopic and ordinal features. To quantify the contribution of different factors to each neuron’s response, we modelled the firing rates (0–1001ms and 100-250ms post-stimulus) of each neuron across word stimuli as a linear combination of word position, retinotopic letter position, and ordinal letter position factors (similar to the linear model used by Agrawal & Dehaene, 2025, to model fMRI and MEG signals; see methods; Figure 4B). Given that the independent variables were correlated, we employed cross-validated LASSO regression to estimate sparse solutions, allowing us to identify the most significant predictors for each neuron.

**Figure 4:**
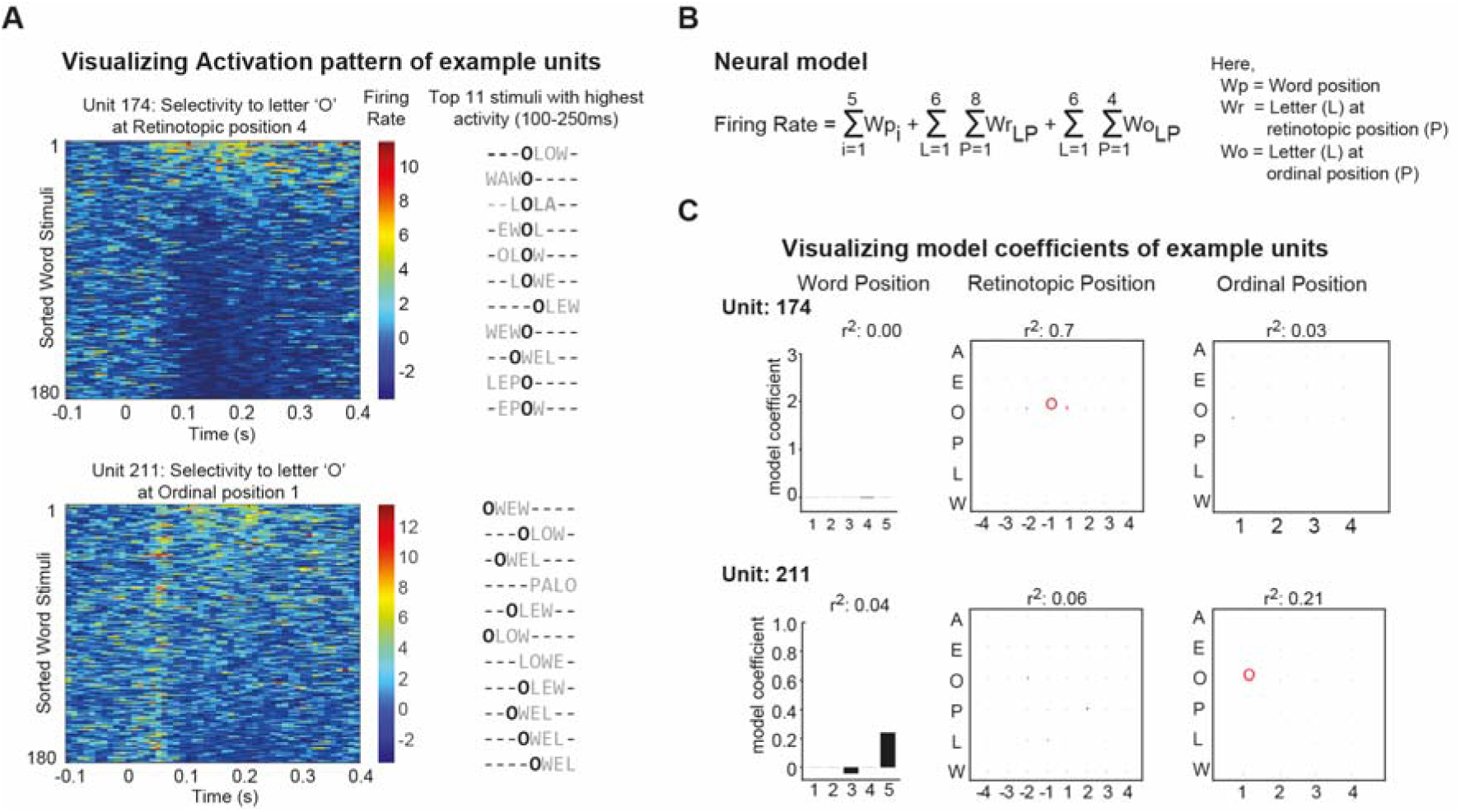
Response profile of two example units. (A) Heatmaps displaying the normalized firing rates of each neuron across all word stimuli. The y-axis represents individual word stimuli, sorted based on their mean firing rate, and the x-axis indicates time after stimulus onset (-100 to 4001ms). Warmer colors denote higher firing rates (left). The top 11 stimuli that elicited the highest firing rates for each neuron are displayed. The letters of interest are highlighted to visualize the property of given units (right). (B) Equation of the linear model that is used to predict the neural responses as a linear combination of word position, retinotopic and ordinal letter positions. (C) Model coefficients from the LASSO regression along with the variance explained for unit 174 (retinotopic) and 211 (ordinal), highlighting the factors contributing to their responses.

The model coefficients reliably predicted the tuning properties of individual neurons. For example, unit 174 showed the highest activity in response to stimuli containing the letter ‘O’ at the fourth position relative to the screen (i.e. just left of fixation). Correspondingly, the model assigned the highest coefficient to the retinotopic representation of ‘O’ at that position, while all coefficients related to ordinal positions were near zero, indicating pure retinotopic encoding (Figure 4C). In contrast, unit 211 responded most strongly to stimuli containing ‘O’ at the first position relative to the word (i.e., the initial letter), indicating an ordinal code.

To quantify the contribution of each factor in predicting the variability in firing rates, we performed a hierarchical stepwise regression analysis. Starting with word position terms, we sequentially included retinotopic terms and then ordinal terms, calculating the increase in explained variance (Δr²) at each step. This analysis confirmed that unit 174 was primarily a retinotopic neuron, as the inclusion of retinotopic terms significantly increased the explained variance (Δr²1=70**%**, p1<10.01), while adding ordinal terms had marginal effect (additional Δr²1=3%, p1<10.05). Conversely, neuron 211 showed a substantial increase in explained variance upon adding ordinal terms (additional Δr²1=21%, p1<10.01) along with the retinotopic terms (Δr²1=6%, p1<10.01), indicating its primary reliance on ordinal position encoding.

We repeated this analysis across all neurons with significant split-half consistency in the IT cortex and found that word position and retinotopic factors explained a larger proportion of variance in neural responses compared to ordinal coding factors. During the late phase (100–250 ms post-stimulus), neuronal encoding preferences shifted from word position toward retinotopic and ordinal position coding (Figure 5A). Post-training, overall model fits increased, indicating enhanced encoding of stimulus features. In particular, ordinal position coding increased significantly during the late time window for monkey 1 (Figure 5A). In monkey 2, the effect of training was largely restricted to retinotopic and word position coding (Figure 5B). Similar trends were observed when model fits were compared across all neurons (Figure S11).

**Figure 5:**
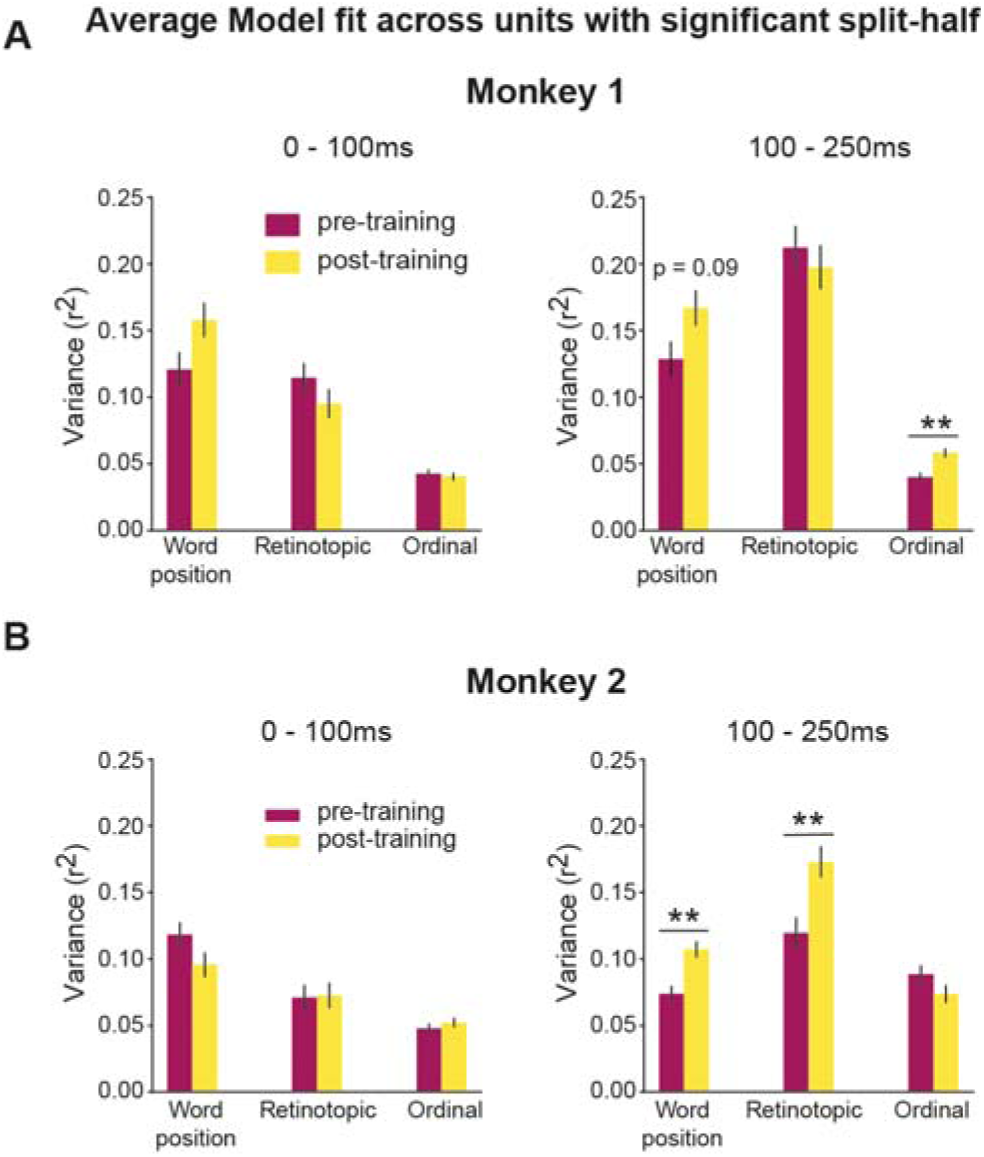
Model fit performance across neurons. (A) Proportion of variance explained by word position, retinotopic, and ordinal factors during early (0–1001ms) and late (100–2501ms) time windows, averaged across all units with good split-half consistency. Asterisks indicate statistical significant difference between pre- and post-training variance (**, p< 0.005 paired t-test) (B) Same as (A) but for Monkey 2

Since our wireless recording setup allowed us record from the same set of neurons over multiple days, it enabled us to examine how tuning evolved within the same neuronal population. Neurons that were strongly modulated by ordinal or retinotopic factors maintained their tuning after training (Figure S11-S12). This result is similar to the children fMRI finding that, as some voxels become tuned to written words (VWFA) during reading acquisition, they maintain their prior tuning to objects (Dehaene-Lambertz et al., 2018). It suggests that literacy training primarily reorients the tuning of neurons that are initially less specialized.

To test whether the increase in ordinal-position coding in the late window of monkey 1 could be explained by greater or faster eye movements after training (Figure S5), we compared receptive-field estimates for a retinotopic unit (unit 174) across spatial positions. Receptive fields remained largely unchanged in the chosen time windows (up to 250ms), indicating stable spatial tuning (Figure S13). Specifically, we trained linear models independently on stimuli presented at each word position and confirmed that the estimated receptive fields were nearly identical across time windows. As expected, explained variance was much lower for words presented in the ipsilateral visual field (r² = 0.04), ruling out eye-movement artifacts. Moreover, monkey 2 showed comparable eye movements but no corresponding increase in ordinal-coding variance post-training. These results indicate that the enhanced ordinal representation observed in monkey 1 reflects genuine changes in neural encoding rather than oculomotor confounds.

### Simulations Using Convolutional Neural Networks (Cornet Z)

To simulate how word representations evolve with training, we employed a convolutional neural network (CNN), specifically Cornet Z (Figure 6A), following the same architecture and training protocol that we previously used to simulate the acquisition of human literacy (Agrawal & Dehaene, 2024; Hannagan et al., 2021). The network was initially pre-trained on object recognition tasks before being recycled for word recognition, in agreement with the neuronal recycling hypothesis (see methods). Once trained, we analyzed the activations across layers in the network corresponding to visual areas (V1, V2, V4, IT, and avgIT), focusing on units that showed selectivity for word stimuli. To quantify the contribution of word position, retinotopic position, and ordinal letter position to the network’s responses, we used the same LASSO regression model that was used in the monkey experiments to fit the activations of word-selective units across the 180 word stimuli used in this study (36 words X 5 positions).

**Figure 6:**
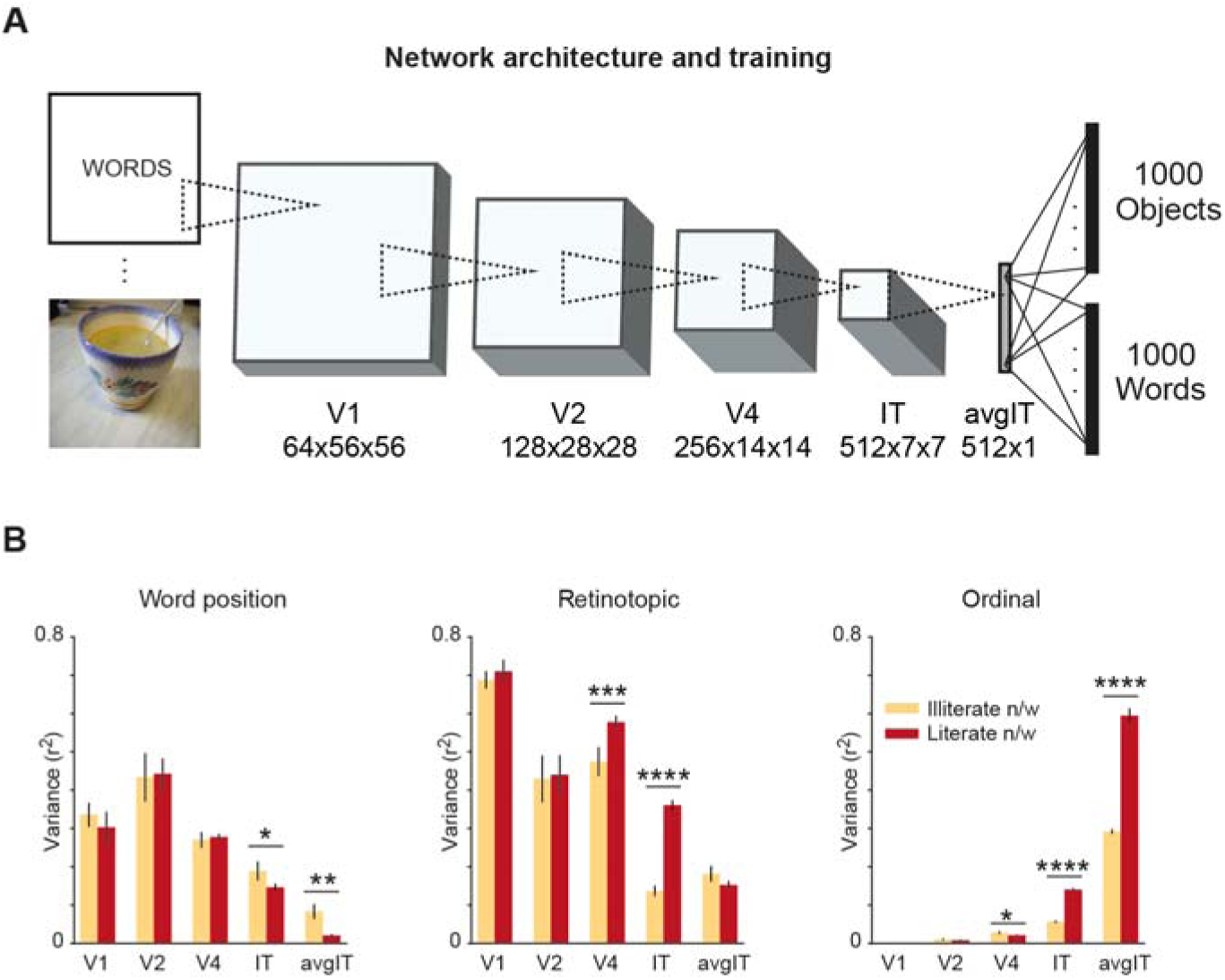
Neural codes observed in simulations of a CNN model of reading. (A) Architecture of the convolution neural network (Cornet-Z) used to simulate the orthographic training. This network was initially trained to only categorize ImageNet dataset, which contains images of objects from 1000 categories. Later, it was jointly trained on both ImageNet and words dataset (B) Proportion of variance explained by different factors of the encoding model for both literate (red) and illiterate network (orange). Asterisks indicate statistical significance (*, p < 0.05; **, p < 0.005 paired t-test)

The results revealed a clear progression of coding across the network layers. In the early layers (V1, V2), unit activity was primarily explained by word position and retinotopic letter position, similar to the encoding observed in early visual cortex. However, as information flowed through later layers (V4, IT, and the higher layers), the contribution of word position and retinotopic encoding progressively decreased, while the proportion of variance explained by ordinal letter position steadily increased (Figure 6B). This shift toward ordinal coding in the deeper layers closely mirrors the neuronal dynamics observed in the monkey IT cortex, where early time windows preferentially encoded word position followed by ordinal encoding. These results suggest that both artificial and biological systems develop more abstract representations of ordinal letter position as information is processed through higher-order visual areas. Interestingly, we observed that even in an “illiterate” version of the network, trained exclusively on object recognition without any exposure to written words, there was evidence of ordinal coding, albeit at a much weaker level (Figure 6B). This finding, is similar to our findings of ordinal units in monkeys on the first day of recording. It suggests that the visual system may possess an inherent predisposition for processing ordinal information, which is then refined through training.

### Test of competing bigram models of word recognition

To evaluate whether IT responses could be better explained by other prominent models of reading rather than by single-letter features, we compared the performance of the letter × position model with two variants of bigram models: (1) a position-dependent bigram model based on the CV and VC letter pairs used to construct the word stimuli, and (2) an open bigram model that captured all ordered letter pairs irrespective of their spatial separation. Like the letter-position model, the bigram models were augmented with the word-position terms (5 regressors) to capture the major effect of retinotopic word position on firing rates and avoid differences driven by positional rather than bigram-specific features (see Methods). Model fits were computed for neurons with significant split-half reliability using 5-fold cross-validated LASSO regression during the early (0–100 ms) and late (100–250 ms) time windows, as described for the letter model.

Across both monkeys, the letter × position model consistently explained the largest proportion of variance, significantly outperforming both bigram models during both pre- and post-training sessions (Figure S15). Similar trends were observed across all time windows with relatively higher model fits during the late time window. Together, these results indicate that IT responses to letter strings were dominated by position-specific letter coding, with only limited contribution from bigram-level representations.

## Discussion

This work provides novel insights into the single-neuron mechanisms underlying the acquisition of orthographic processing in non-human primates. Individual neurons in the inferior temporal (IT) cortex of macaque monkeys were found to encode both retinotopic and ordinal letter positions within words. Notably, ordinal encoding units were present even before training. With training, the neural representation space evolved in direct relation to behavior. Training on orthographic tasks enhanced the ordinal representation in monkey 1, whose performance improved in a location-invariant match-to-sample task, but it only improved the retinotopic representation in monkey 2, who did not improve. These findings suggest that training refines pre-existing neural circuits to facilitate efficient orthographic processing. Simulations of orthographic learning in convolutional neural networks (CNNs) mirrored these observations, showing increased ordinal encoding in deeper layers of the network.

### Written word recognition as cortical recycling

To our knowledge, this is the first study to directly compare neuronal responses to written letter strings in non-human primates over a specified training interval, bridging a critical gap in understanding how the primate visual system adapts to support culturally acquired skills like reading. The results provide empirical support for the neuronal recycling hypothesis (Dehaene & Cohen, 2007), which posits that cultural inventions like reading co-opt pre-existing neural circuits evolved for different but similar functions. Two aspects of the results fit with this hypothesis: (1) The presence of neural preferences for specific letters even in untrained monkeys, as predicted if the shapes that we selected as letters were chosen because they were already easily encoded by the preliterate visual system (Changizi et al., 2006; Dehaene, 2009; Szwed et al., 2009); (2) the presence of an ordinal code prior to training, and its enhancement after training, compatible with the idea that the visual system repurposes existing neural architectures to accommodate new cognitive demands. The presence of ordinal encoding units prior to training indicates that the IT cortex has inherent capabilities for processing sequential visual information, which can be fine-tuned through experience.

Recent simulations and human brain-imaging studies have further elaborated on this hypothesis, suggesting that the ventral visual pathway is particularly amenable to such recycling due to its hierarchical organization and plasticity (Agrawal & Dehaene, 2024, 2025; Dehaene et al., 2015; Hannagan et al., 2021). The fact that, in humans, written words activate a reproducible cortical site (the VWFA in the left occipito-temporal sulcus) seems to be due to a combination of two factors: the specific visual features of words, which put them in the category of high-resolution foveal (Hasson et al., 2002) and “spiky” shapes (Bao et al., 2020), and the pre-existing connectivity of the human left occipito-temporal cortex with left-hemispheric temporal and inferior frontal language areas that are the targets of fluent reading (Barttfeld et al., 2018; Hannagan et al., 2015; Pinel & Dehaene, 2009; Saygin et al., 2016). Here, however, we trained adult monkeys. Since they lack such language areas, and since limits on adult plasticity may prevent the self-organized grouping of visual stimuli with similar features at a specific cortical sites, unlike in juvenile monkeys (Srihasam et al., 2012), we expected that orthographic codes would be broadly distributed in inferotemporal cortex, especially after a short training period (5 days). This may explain that letter-encoding neurons could be observed in the present study even in the absence of prior fMRI-based identification of relevant recording sites.

### Neural code for written words

The present work clarifies the neural code by which the ventral visual pathways achieve an invariant recognition of letter strings, a hotly debated issue in the cognitive neuroscience of reading (Dehaene et al., 2005; Grainger & Van Heuven, 2004; McCloskey et al., 2013; Norris, 2013; Whitney, 2001). The challenge is the written word recognition system must efficiently recognize visual forms that are visually quite similar (here, 4-letter words) and made of the same parts (here, 6 letters), yet do so invariantly across changes in word size and retinotopic position. Neuropsychological studies (McCloskey et al., 2013), simulations (Agrawal & Dehaene, 2024; Hannagan et al., 2021) and human behavioral (Agrawal et al., 2019) and brain-imaging (Agrawal & Dehaene, 2025) increasingly hinted a possible solution: neurons tuned to both specific letters and their ordinal position, relative either to the left or to the right end of the word. The present study is the first to obtain direct evidence for the existence of such neurons in monkey IT and their amplification by orthographic learning: neurons sensitive primarily to the first letter of a 4-letter word were found in right-hemispheric IT, and neurons sensitive to the last letter in left-hemispheric IT. There were very few neurons tuned to inner letter positions (2^nd^ or 3^rd^), like in the model (Agrawal & Dehaene, 2024). Furthermore, monkeys showed no evidence of a sensitivity to the transposition of inner letters, an effect which emerges with reading expertise in humans (Hasenäcker & Schroeder, 2022; Ziegler et al., 2014).

How do IT neurons acquire their sensitivity to ordinal position, irrespective of the word’s retinotopic location? Simulations suggest that ordinal selectivity emerges by pooling such retinotopic units as well as detecting the presence of a space left and right of each word, thus allowing neurons to detect the location of letters relative to that space (“space bigram” units; Agrawal & Dehaene, 2024). The present work confirms that, at the same cortical location, many units are sensitive to retinotopic letter location, but much denser recordings would be needed to test the model’s hypothesis that the receptive field of ordinal-position units arises from inhibitory and excitatory letter-tuned retinotopic units (Agrawal & Dehaene, 2024).

### Role of the Inferior Temporal Cortex in Orthographic Processing

The present findings contribute to a growing body of research highlighting the crucial role of the IT cortex in visual word recognition and orthographic processing. Katti and Arun (2022) demonstrated that a separable neural code in monkey IT cortex enables decoding of complex visual stimuli such as CAPTCHAs, suggesting a rich and adaptable neural code where responses to n-grams can be predicted by a sum of responses individual letters. The current study aligns with this perspective, but further shows that IT neurons do not merely implement a “bag of letter” representation of words, but encode individual letters at specific ordinal positions. While the CAPTCHA study primarily focussed on decoding the accuracy of letter identity or positions, we use encoding models to identify units that represent letter identity and ordinal letter positions. The specificity of IT units to certain letters was also observed in earlier studies that investigated the full set of 26 characters (Rajalingham et al., 2020). These units, recorded from the IT cortex of untrained monkeys, implemented neural representations that could predict the error patterns in word recognition tasks observed in baboons and humans (Grainger et al., 2012), suggesting that the non-human primate provides an adequate model for a cortical precursor of orthographic processing. However, those previous studies did not explore the effect of training on visual representations of words. In humans, behavioral and brain imaging data suggest that learning to read enhances the compositionality of word representation: with literacy, word responses become more predictable from single letter responses (Agrawal et al., 2019, 2020). Although the primates in our study were trained on orthographic tasks for limited days, the findings nicely align with the evidence from human experiments, supporting the idea that reading acquisition refines pre-existing neural circuits and enhances ordinal coding.

### Neural Plasticity and Training Effects

Our results shed light on the neural plasticity associated with orthographic training. While several previous studies on perceptual learning have reported decreases in neural activity following training (Koyano et al., 2023; Meyer et al., 2014; Woloszyn & Sheinberg, 2012), here an overall increase in mean neural activity was observed post-training. This divergence may be attributed to differences in the nature of the stimuli. Most studies on image familiarity recruit stimuli that preserve the natural statistics of the monkey’s environment, such as unfamiliar faces and objects. However, word stimuli have distinctive properties; letters are more similar to each other and have higher spatial frequencies than objects. During the initial stages of learning, additional neural resources probably need to be recruited to encode and process those complex and unfamiliar stimuli. Indeed, we found a greater differentiation of IT neural responses to different letter strings after training (Figure 3C), in agreement with the hypothesis that literacy leads to an expanded neural representational space, with more distinct and therefore more efficient neural representations (Agrawal et al., 2019; Meyer et al., 2014). The observation that visually similar letters are closer to each other in neural space (Figure 3D) is also consistent with the visual representation space of literate humans (Agrawal et al., 2020).

While it is tempting to associate the increase in IT neural activity with the formation of a Visual Word Form Area (VWFA), whose activation to letter strings also increases sharply during reading acquisition (Brem et al., 2010; Dehaene-Lambertz et al., 2018; Feng et al., 2022), caution is warranted. First, our study was performed in adult monkeys, but a prior fMRI study of training with letter and number symbols found the emergence of a dedicated IT subregion similar to the VWFA only in juvenile but not in adult monkeys (Srihasam et al., 2012); our experiment might have yielded different results, and might also have been more pertinent to human reading acquisition, if it had been performed in juvenile animals. Second, in humans, the VWFA may form specifically when letter-sound correspondences are learned (Brem et al., 2010), while we focused solely on visual orthographic learning. Third, intriguingly, we observed a greater neural response to strings of unfamiliar letters in monkey IT cortex, whereas the VWFA in literate humans shows a reduced response to false fonts and infrequent letters (Vinckier et al., 2007; Woolnough et al., 2021; Zhan et al., 2023). In this respect, our results are more similar with human fMRI findings of higher responses to novel untrained scripts in the lateral occipital cortex (Agrawal et al., 2019).

### Limited Impact of Eye Movements

Although stimuli were presented briefly to minimize eye movements, small saccades were observed, particularly when stimuli were presented at peripheral positions. Could ordinal position coding be partly attributed to these eye movements? Several observations argue against this possibility. First, the amplitude of eye movements, when they occurred, did not exceed a single letter. The observed ordinal position coding was consistent across a range of stimulus positions, not just those affected by eye movements. Second, they came after ∼200 ms, but the ordinal code was observed prior to that point. Third, retinotopic units exhibited stable and sharp tuning up to 250 ms, suggesting that any shift in the retinal image caused by saccades remained very small. Previous studies have shown that IT neurons can maintain object selectivity despite changes in retinal position—a phenomenon known as position tolerance (Hung et al., 2005)—supporting the robustness of these findings.

### Implications for Reading Disorders

The observed neural code has implications for understanding reading errors in normal subjects as well as dyslexia and other reading disorders, which may result, at least in part, from atypical development or functioning of the neural circuits involved in orthographic coding (Feng et al., 2020; Goswami, 2015). It was previously argued that the properties of the IT cortex may explain common errors in letter recognition among children, such as left-right inversion errors, which have been linked to the tendency of IT neurons to respond similarly to mirror images of objects (Dehaene, Nakamura, et al., 2010). Here, behavioral and neural converged to indicate that monkeys preferentially encoded the first and last letters of a word, but had greater difficulty encoding the specific position of inner letters, therefore tending to respond “same” to test items where the inner letters were changed or swapped. Such an imprecision in the code for internal letters may explain letter transposition effects in normal reading, whereby orthographic transpositions such as “bagde” may not be detected and may prime the original word (badge) (Perea & Lupker, 2003; Schoonbaert & Grainger, 2004; Ziegler et al., 2013). If internal positions became very imprecise, it could also explain the developmental deficit known as “letter position dyslexia”, whereby children fail to precisely encode the position of inner letters and misread for instance “form” as “from” (Friedmann & Gvion, 2001; Potier Watkins et al., 2023). In both normals and letter-position dyslexics, such errors spare the first and last letter, in full agreement with their sharp positional encoding in the present work. Interventions targeting the enhancement of ordinal encoding mechanisms could potentially ameliorate those difficulties.

A distinct subtype of dyslexia, attentional dyslexia (Friedmann et al., 2010; Potier Watkins et al., 2023; Warrington et al., 1993), involves swapping the letters at the same position in two different words, often the first or the last position, for instance misreading “win fed” as “fin wed”. Again, similar errors also occur in normal readers under conditions of fast presentation and divided attention (Treisman & Souther, 1986). This error pattern could be explained by neurons encoding ordinal letter positions, if they did so for two words at once (as suggested by the fact that they respond identically regardless of retinotopic word position). Here, interventions could target the focusing of attentional and processing resources on a single word at a time. More broadly, understanding how training refines neural circuits may thus inform educational strategies and therapeutic interventions for reading disorders.

### Limitations and Future Directions

A key limitation of this study is its relatively short training period (5 days), which may not fully capture the extensive process of literacy acquisition in humans. More extended training might allow the monkeys to develop a more precise neural code for inner letters, as well as more nuanced discriminations between different types of nonwords, as observed in human readers who show graded neural responses to word-likeness (Binder et al., 2006; Vinckier et al., 2007; Woolnough et al., 2021; Zhan et al., 2023). Future studies could involve longer training durations, but also tasks that incorporate phonological and semantic integration, to more closely mimic human reading acquisition. Additionally, incorporating other modalities, such as auditory stimuli, could provide insights into how multimodal integration influences orthographic processing.

While CNN simulations paralleled empirical findings, the model remains limited in many ways. It is purely feedforward, while feedback connectivity clearly plays a role in human and presumably monkey responses to words (Woolnough et al., 2021). Also, the network was trained on word recognition using cross-entropy loss, not the specific delayed match-to-sample task performed by the monkeys. Training the network under conditions that more closely mimic the monkeys’ experience could enhance the relevance of the model to these findings. Recent advancements in deep learning architectures that incorporate feedback connectivity, attention mechanisms and sequential processing could provide more accurate models of neural processing during reading tasks (Hannagan et al., 2021; Yamins & DiCarlo, 2016).

## Methods

All experimental procedures were approved by the Institutional Animal Ethics Committee of the Indian Institute of Science (CAF/Ethics/399/2014 and CAF/Ethics/750/2020) and by the Committee for the Purpose of Control and Supervision of Experiments on Animals, Government of India (25/61/2015-CPCSEA and V-11011(3)/15/2020-CPCSEA-DADF).

### Animals

Two adult male bonnet macaque monkeys (*Macaca radiata*), designated M1 (Di) and M2 (Co), aged approximately 9 years and weighing 6.61kg and 51kg respectively, participated in this study. The monkeys were housed in play area in a temperature-controlled environment with a 12-hour light/dark cycle. All training and experimental sessions were conducted during the light phase.

### Experiment design

The study spanned seven consecutive days for each monkey. On day 1, the monkeys performed an RSVP fixation task (pre-training). This was followed by five days (days 2–6) of training on a delayed match-to-sample (DMS) task. On day 7, the monkeys performed the RSVP fixation task again (post-training) to evaluate changes in neural representations due to training. Tasks were conducted at the same time each day to control for circadian influences. Both monkeys had extensive prior experience with the match-to-sample paradigm using a large range of non-orthographic visual stimuli (natural images). Therefore, during the study, we expected them to focus on learning the properties of new letter string stimuli rather than on acquiring the task structure itself.

### Behavioral Tasks

#### RSVP fixation tas

Each trial began with a hold cue displayed on the right side of a touch screen monitor (Figure11B). The monkey initiated the trial by touching the hold cue with its hand. A fixation cross then appeared at the center of the screen, and the monkey was required to maintain its hand on the hold cue and fixate within an 8° radius around the fixation cross. For rapid serial visual presentation (RSVP), a letter-string stimulus was presented for 200 ms, followed by a 200 ms blank screen. This sequence was repeated eight times per trial using distinct unrelated stimuli. Successful maintenance of fixation throughout the trial was rewarded with juice. A total of 396 correct fixation trials were conducted per session.

#### Delayed match-to-sample (DMS) task

Trials began similarly with the monkey touching the hold cue to initiate the trial (Figure11B). After fixating on the central cross, a sample stimulus appeared for 200 ms, followed by a 200 ms blank screen. Subsequently, the test stimulus appeared at the center, the hold cue disappeared, fixation constraints were lifted, and two choice buttons appeared above and below the previous location of the hold cue. The monkey had up to 5 s to respond by touching one of the choice buttons: the upper button if the test stimulus matched the sample, or the lower button if it was different. Correct responses were rewarded with juice. The test stimulus and choice buttons remained on the screen until the monkey responded or the 5 s elapsed. On each day, a session comprised at least 10 blocks, with each block containing 18 “same” and 18 “different” trials, resulting in a total of 360 trials. Incorrectly responded trials were not repeated during the block. The monkey also made fixations and hold errors in some blocks apart from correct and incorrect responses. Therefore, we repeated those blocks again to ensure that the monkey saw all the 360 stimulus conditions.

### Stimuli

Six visually distinct letters—three consonants (L, P, W) and three vowels (A, E, O)—were used to create four-letter strings. Using those letters, we created 36 stimuli designated as “words”. All words had alternating consonant-vowel structures, either consonant-vowel-consonant-vowel (CVCV) or vowel-consonant-vowel-consonant (VCVC) patterns. We selected 36 unique words (18 CVCV and 18 VCVC) ensuring: 1) equal representation of each letter across all four positions. 2) Balanced occurrence of each consonant-vowel bigram at word beginning and word ending. A list of all 36 words is shown in the table below.

|  |  |  |  |
| --- | --- | --- | --- |
| WAWO | WAPE | AWEP | AWAW |
| WEWE | WEWO | APOP | APAL |
| WOLE | WOPA | ALAW | ALAP |
| PALO | PALE | EWOL | EWEL |
| PEPO | PELA | EPOW | EPOL |
| POPE | POWA | ELAP | ELEP |
| LAPA | LALO | OWEL | OWEW |
| LEWA | LEPO | OPAL | OPOP |
| LOLA | LOWE | OLEW | OLOW |

To assess position invariance and distinguish retinotopic from word-centered encoding, each word was presented at five horizontal positions relative to the central fixation point: -2, -1, 0, +1, and +2 letter widths (with negative values indicating leftward positions and positive values rightward). Positions -2 and +2 corresponded to unilateral hemifield presentations, while intermediate positions included letters in both hemifields and tested shifts in receptive fields. This resulted in a total of 180 word stimuli (36 words × 5 positions).

All other combinations of letters were categorized as nonwords. They were generated to create varying levels of non-match difficulty in the delayed match-to-sample task. For each of the 36 possible sample word, eight types of non-match stimuli were constructed:

- *Unfamiliar letters*: Strings composed of four out of six unfamiliar letters not used to compose words (e.g., G, H, X, I, U, Y), serving as an easy non-match condition.
- *Uniform Letter Strings*: Random strings comprising either only consonants (e.g., PLWP) or only vowels (e.g., AOEO).
- *Orthogonal manipulations of structure and Lexicality*: Four categories of stimuli forming a 2×2 design varying lexicality (word [different from the sample word] or nonword) and consonant-vowel structure (opposite or same as the sample word), probing whether monkeys utilized these features in their judgments. In this part of design, the non-words followed either the same CV structure as the preceding sample word (both CVCV or both VCVC), or the opposite structure (CVCV word followed by VCVC word or pseudoword, or VCVC word followed by CVCV word or pseudoword).
- *Letter Position Errors*:

o Transpositions: Pseudoword consisting in swapping the two internal letters of the sample word.
o Substitutions: Pseudoword consisting in replacing the two middle letters of the sample word with different letters.

As an example, the full list of stimuli for training day 1 is shown in Table S1.

### Stimulus Presentation

#### RSVP Fixation task

In this part of the experiment, the same set of stimuli was used during pre-training and post-training sessions to assess changes in neural representations independent of task performance. A total of 396 stimuli were presented: 180 words (36 x 5 positions) and 216 nonwords (36 x 6 categories, excluding words).

#### DMS task

Word stimuli remained constant across training days, while nonword stimuli were generated anew each day to prevent memorization. The nonwords were also different from the stimuli used in the fixation task. Each day, a list of 180 match and 180 non-match trials were run. To achieve such as balance, we chose only five out of eight possible non-match types for a given word.

Each non-match was randomly chosen and paired with one of the 5 possible positions of a given sample word. Our uniform sampling approach ensured that the occurrence of each non-match type in this task was approximately equal within a given day.

All stimuli were displayed in Consolas, a monospaced font, on a touch screen monitor (34.41cm width × 19.351cm height, resolution 13661×1768 pixels, 601Hz refresh rate). The viewing distance was approximately 23 cm, resulting in stimuli subtending 16 degrees of visual angle.

### Data acquisition

We recorded neural activity wirelessly using chronically implanted microelectrode arrays into the central portion of the inferior temporal cortex (M1: right hemisphere, M2: left hemisphere) as well as premotor/prefrontal regions (M1: right ventral premotor cortex; M2: left ventral premotor cortex & ventrolateral prefrontal cortex; data not included in this study).

Electrode implantation surgeries were performed under sterile conditions under anesthesia. Post-operative care included analgesics and antibiotics as needed. Neural signals were digitized at 25 kHz. We used a combination of high-pass and low pass filters to obtain the spikes from the raw data - a high pass 4^th^ order Butterworth filter (cutoff frequency 250 Hz) and a low pass 4^th^ order Butterworth filter (cutoff frequency 3 kHz). To calculate the threshold for the spike, we first obtained the noise estimate as 75^th^ percentile of the absolute amplitude of filtered signal and then the spike detection thresholds were generally set at 3 times of noise estimate. We changed some of these parameters when necessary while performing spike-sorting. To track the same neurons across the two RSVP fixation sessions (before and after training), we kept waveform detection parameters the same for each neural site. Units that showed identical waveforms in the two RSVP sessions were assumed to be identical. This resulted in 84 single-unit (81 post-training) and 98 multi-unit recordings (98 post-training) in monkey 1, and 75 single-unit (80 post-training) and 119 multi-unit recordings (120 post-training) in monkey 2. We observed similar trends in single and multi-unit activity, so the results are presented using the combined data.

### Data preprocessing

For each neuron, firing rates were calculated by counting spikes within specified time windows. This estimation was restricted to repetitions where the monkey maintained its fixation. To assess temporal dynamics across the stimulus presentation, we performed a time-course analysis using overlapping 20 ms bins with a 10 ms step size. Additionally, and given the presence of two consecutive peaks in average activity profiles, firing rates were averaged over early (0–100 ms) and late (100–250 ms) post-stimulus time windows to examine temporal variations in neural responses. Normalization was performed using z-scores to standardize the firing rates across neurons and sessions. Specifically, we calculated the mean (μ_baseline_) and standard deviation (σ_baseline_) of firing rates during the pre-stimulus window (−1001ms to 01ms) across all stimuli. Post-stimulus firing rates (FR_post_) were then normalized using the equation:

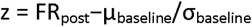

This z-score transformation allowed for the comparison of neural activity across different neurons by accounting for baseline firing rate differences.

#### Behavioral Data

Response times (RTs) were measured from the onset of the test stimulus to the monkey’s response. Trials with RTs exceeding 5s or with premature responses were excluded. Behavioral analyses were performed using Python with the scikit-learn library for statistical modelling.

### Eye Tracking

Eye data was collected using ETL 300HD, a binocular eye-tracker camera with an IR illuminator (ISCAN Inc) at sampling rate of 120 Hz and then sent to the main server. For eye calibration, a separate block was performed on the touchscreen at the beginning of each task (or experiment). The calibration block was initiated when the monkey touched a green circular button of radius 4 dva placed at 20 dva right from centre of the screen. Then the monkey had to maintain gaze at the centre of the screen for 500 ms with a fixation radius of 8 dva. On successful gaze, four grey dots appeared sequentially at the four corners of the screen, and the monkey had to saccade to and maintain the gaze at each location for 300 ms. The fixation radius was set to be 8 or 10 dva around each grey dot. The monkey was rewarded with juice on successful fixation at all the four locations and this was considered as correct trial. We recorded at least two correct calibration trials before moving on to the main task.

### Modelling behaviour data

We used logistic regression to model the monkeys’ binary choices (same or different) on each trial of the delayed match-to-sample task. Data from all five training days were pooled to enhance statistical power. A five-fold cross-validation approach was used to prevent overfitting. The predictor variables included:

1. **Sample Position**: Five binary variables indicating the sample’s position (-2, -1, 0, +1, +2).
2. **Fraction of Common Letters**: Calculated for retinotopic positions, ranging from 0 to 1 in increments of 0.25.
3. **Identical letters at ordinal positions**: Independent binary indicator for each of the 4 ordinal positions, indicating whether a letter in the sample stimulus matched the letters in the test stimulus at a given ordinal position.
4. **Match Condition**: Binary indicator of whether the full 4-letter test stimulus matched the sample word (1 for match, 0 for non-match).
5. **Test Stimulus Lexicality**: Binary indicator of whether the test stimulus was a word (excluding exact matches).
6. **Unfamiliar letters**: Binary indicator for test stimuli composed of unfamiliar letters.
7. **Word Structure Match**: Binary indicator for test stimuli having the same consonant-vowel structure as the sample.
8. **Training Day**: Five binary variables representing each training day.

In the logistic regression model, Y = Xb, Y represents the binary response variable of size (ntrials x 1), X is a predictor-variables matrix of size (ntrials x 19), b is a unknown coefficients vector of size (19 x 1). To assess the variability of model coefficients, we performed bootstrap resampling with replacement (n = 100 iterations). Response time data were analyzed using cross-validated LASSO regression due to the continuous nature of RTs and to perform feature selection.

### Modelling neural responses

For each unit, we modelled its variation in firing rate across 180 word stimuli as a linear combination of the following three stimulus parameter types, as described in figure 4B:

1. **Word Position**: This factor captured the spatial position of the entire word stimulus relative to fixation. We used a one-hot encoding scheme to generate a matrix representing the five possible word positions for each of the 180 stimuli. The resulting matrix had dimensions of 180 (stimuli) by 5 (word positions).
2. **Retinotopic Position**: This factor represented the presence of each letter at specific retinotopic locations on the screen. For each of the eight possible retinotopic positions (four positions left and four positions right of fixation), binary vectors were created to indicate the presence or absence of each of the six possible letters. The final matrix had dimensions of 180 (stimuli) by 48 features (6 letters × 8 positions).
3. **Ordinal Position**: This factor encoded the identity of letters at each ordinal position within the word (first letter, second letter, etc.). Similar to the retinotopic model, we used a one-hot encoding for each of the six possible letters and four ordinal positions, resulting in a matrix of 180 (stimuli) by 24 features (6 letters x 4 positions).

Thus, in the linear model Y = Xb; Y is a vector of size (180 x 1) indicating the firing rate of a given unit; X is a matrix of size (180 x 77) excluding the constant term; and b is an unknown weight vector of size (77 x 1). Model coefficients were estimated using 5-fold cross-validated LASSO regularized linear regression (LassoCV function in python).

### Bigram model

In addition to the letter-based models, we implemented two bigram-based models to test whether IT neuronal responses were better explained by combinations of adjacent letter pairs rather than by single-letter features. Both models were constructed using the same set of 180 word stimuli and were designed to capture different structural aspects of letter co-occurrence patterns.

1. *Position-Constrained Bigram Model:* Given that our stimuli followed a fixed CVCV or VCVC structure, we restricted the bigrams to all possible CV letter pairs (n = 9) and VC letter pairs (n = 9). bigrams were further defined according to their within-word position. There were 18 possible bigrams in the first half of the word (positions 1–2) and 18 in the second half (positions 3–4), yielding a design matrix of 180 (stimuli) × 36 (bigrams).
2. *Open bigram model:* This factor encoded the presence of all possible ordered pairs of letters (bigrams) within each word stimuli, regardless of their spatial separation, provided that the first letter preceded the second. For the six possible letters used in our word set, all pairwise combination resulted in 36 bigrams. The resulting design matrix had dimensions of 180 (stimuli) × 36 (bigrams).

Additionally, word position terms (n = 5) were included for equitable comparison with the letter x position model. As before, the relationship between neural responses and these bigram features was expressed as **Y=Xb**, where Y is the vector of firing rates (180 × 1), X is the corresponding bigram feature matrix (180 x 41), and b is the vector of regression weights. Model coefficients were estimated using 5-fold cross-validated LASSO-regularized linear regression (LassoCV).

### Split-half measures

To evaluate the reliability of the neural responses, we computed split-half consistency for each neuron across all 396 word and nonword stimuli presented during the RSVP fixation task. For each stimulus, we divided the repeated presentations into odd-numbered and even-numbered trials to form two independent response sets. The normalized firing rates for each neuron were computed for each set over the 0–200 ms post-stimulus window, yielding two response vectors of length 396 (one per split). We then computed the Pearson correlation coefficient between these two vectors to quantify the neuron’s response consistency. This correlation served as a measure of the neuron’s reliability across repeated presentations of the full stimulus set. The number of units with significant split-half consistency (p1<10.05) was as follows: for monkey 1, 109 units pre-training and 94 units post-training; for monkey 2, 128 units pre-training and 103 units post-training.

### Neural Network Simulation

To simulate changes in the primate’s neural representation space before and after training, we used convolutional neural networks (CNNs) that were initially trained only on objects (ImageNet) and later trained jointly on images of French words and objects. We employed pre-existing models from earlier studies (Agrawal & Dehaene, 2024, 2025; Hannagan et al., 2021), which had already been analyzed and shown to exhibit properties consistent with evidence from orthographic processing studies. To provide a comprehensive overview, here we reiterate the training protocol from those studies.

In the first phase, the network was trained on the ImageNet dataset, which contains approximately 1.3 million images across 1,000 categories. This network, considered “illiterate,” encoded the visual properties of objects but lacked any representation of text. In the second phase, we extended the number of output nodes to 2,000 (1,000 for images and 1,000 for words), with full connectivity to the avgIT layer units. The entire network was then retrained jointly on the ImageNet dataset and a synthetic word dataset, which also consisted of 1.3 million images of 1,000 different words. After this phase, the network was considered “literate.“

PyTorch libraries was used to train these networks, utilizing stochastic gradient descent with categorical cross-entropy loss. The learning rate, initialized at 0.01, was scheduled to decrease linearly with a step size of 10 and a default gamma value of 0.1. Phase 1 training lasted for 50 epochs, and phase 2 for an additional 30 epochs, after which classification accuracy no longer improved. To enhance performance, ImageNet images were transformed using standard operations such as “RandomResizedCrop” and “Normalize,” with image dimensions set at 224×224x3. In the word dataset, to avoid cropping letters, the default scale parameter of RandomResizedCrop was modified so that 90% of the original image was retained. For fairness, we did not apply operations like flipping to the ImageNet dataset, as the same operations on the word dataset would create mirror images of words, which is not typical in reading.

The French word dataset included frequent words of 3 to 8 letters in length. The synthetic dataset contained 1,300 stimuli per word for training and 50 stimuli per word for testing. Variations in the stimuli were created by altering position (shifted -50 to +50 along the horizontal axis, and -30 to +30 along the vertical axis), size (ranging from 30 to 70 pts), fonts, and case (both English and French). There was a total of five different fonts: Arial and Times New Roman for the training set, and Comic Sans, Courier, and Calibri for the test set.

#### Identification of word-selective units

Similar to fMRI localizer analysis, a unit was considered word-selective if its responses to the trained French words were greater than its responses to non-word categories (faces, houses, bodies, and tools) by at least 3 standard deviations. Body and house images were sourced from the ImageNet dataset, tools from the “ALET” dataset, and face images from the “Caltech Faces 1999” dataset. We randomly selected 400-word stimuli and 100 images from each of the other categories to identify category-selective units.

