## Supplementary for "Orthographic training finetunes a neuronal letter position code in primate visual cortex"

| **Words** | **Unfamiliar Letters** | **All Conson-**  **ant and vowel strings** | **Non-**  **words of different structure** | **Words of different structure** | **Non-**  **words of same structure** | **Words of same structure** | **Letter**  **Trans-position** | **Letter**  **Substi-tution** |
| --- | --- | --- | --- | --- | --- | --- | --- | --- |
| WAWO | YIIY | LLWL | OLEP | EWEL | LALE | WEWE | WWAO | WLLO |
| WAPE | UUIH | WLPL | APOW | EPOL | WALE | LEPO | WPAE | WLEE |
| AWEP | HGGH | AEOO | WAPA | LEWA | ELOW | OLOW | AEWP | AALP |
| AWAW | HGXG | OAAE | WEWA | WEWO | APOW | AWEP | AAWW | AOLW |
| WEWE | IYYY | LWLW | EPAP | ELAP | LEWE | WEWO | WWEE | WLAE |
| WEWO | IYXG | PLPP | ALAL | APOP | LEWO | LOLA | WWEO | WLLO |
| APOP | UYYY | AEEE | WOLO | WEWE | OWOW | OPOP | AOPP | AAAP |
| APAL | GUXU | AAOA | POWE | WAPE | ALOP | OWEL | AAPL | AELL |
| WOLE | UIXU | LLLP | OLAW | APOP | POLE | PALE | WLOE | WPAE |
| WOPA | XUXG | PWWP | ALEL | OPOP | WALO | LEWA | WPOA | WAAA |
| ALAW | UUUU | OOAO | LEWE | LAPA | EWOP | ELEP | AALW | AEEW |
| ALAP | GHXY | OEOE | LOWO | LOLA | APEW | OWEW | AALP | AOEP |
| PALO | YIUX | WLPP | ALAL | OLOW | POPA | WEWE | PLAO | POOO |
| PALE | UGUU | WLPL | APAP | ALAW | WOWA | LALO | PLAE | PEWE |
| EWOL | YUGU | OEOE | WOPE | PELA | ELEW | EWEL | EOWL | EALL |
| EWEL | IGUX | AOOO | WEPE | LEWA | OWAL | AWAW | EEWL | EPPL |
| PEPO | UGGX | WPPW | OWAW | OPAL | WOWE | LEWA | PPEO | PLWO |
| PELA | XIYH | WPWL | OLOP | EWOL | PEPE | POWA | PLEA | POPA |
| EPOW | HYUY | AOEO | LEPE | WEWO | OPOL | OWEL | EOPW | ELAW |
| EPOL | XHYH | OEAA | WOWO | LOLA | EWAW | AWAW | EOPL | EAAL |
| POPE | GXUY | LPPL | EPAW | AWAW | LEPA | WEWE | PPOE | PLEE |
| POWA | XGHH | LPLL | ALEL | AWEP | LEWO | LEPO | PWOA | PAEA |
| ELAP | IIHX | EEOE | WALE | WEWO | AWAP | ALAW | EALP | EPOP |
| ELEP | HGII | AAAE | POLO | LALO | ELOP | OWEL | EELP | EWOP |
| LAPA | XHYU | LPLW | EWAL | ELAP | PEPA | WOLE | LPAA | LLOA |
| LALO | UXII | WPPW | APOL | OWEW | LOPA | LOLA | LLAO | LOEO |
| OWEL | IHGG | EEOA | WOWA | WOLE | EWAP | ALAP | OEWL | OOPL |
| OWEW | UXIH | OEAA | WEPA | POWA | EPOP | AWAW | OEWW | OAPW |
| LEWA | IGYX | LLLW | OWEP | ALAP | PEPA | WAPE | LWEA | LLLA |
| LEPO | UXIY | PWLP | ALAL | ALAP | POWO | LOWE | LPEO | LWAO |
| OPAL | XUGH | AOOA | LEWE | POPE | OPEW | OLEW | OAPL | OWWL |
| OPOP | IXUX | OEOO | LAWE | LOWE | OWAW | APOP | OOPP | OAAP |
| LOLA | IGIY | LPWP | AWOL | OWEL | POWE | WAWO | LLOA | LAEA |
| LOWE | UIYU | PWWW | OPEL | EWEL | PEPE | LAPA | LWOE | LPAE |
| OLEW | IXXY | OOOE | WAWE | POPE | APAP | OPAL | OELW | OAAW |

**Table S1:** A full list of the stimuli used on training day 1. The nonwords were unique on each the five days to avoid memorization.

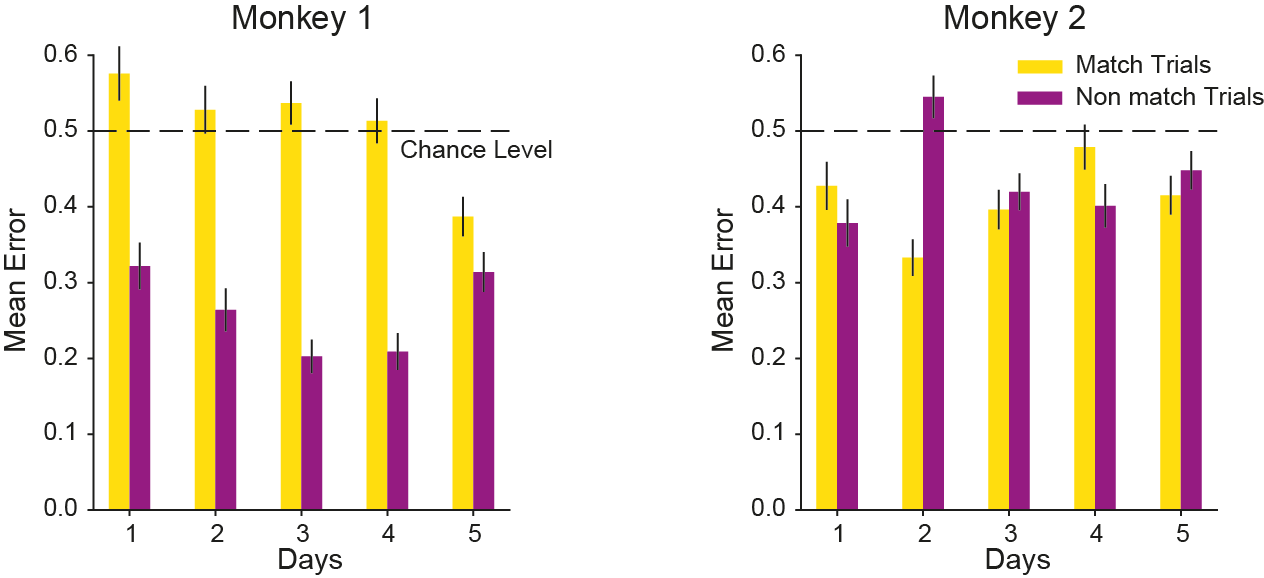

**Figure S1: Behavioral Performance in the match-to-sample task**

**Mean error rate for match and non-match conditions across different days for monkey 1 (left) and monkey 2 (right). Dashed horizontal line indicates chance level. Error bars indicate standard deviation across 100 bootstrap estimates. The better performance of monkey 1 was mainly driven by non-match conditions and it learnt to accurately respond to match conditions only on day 5.**

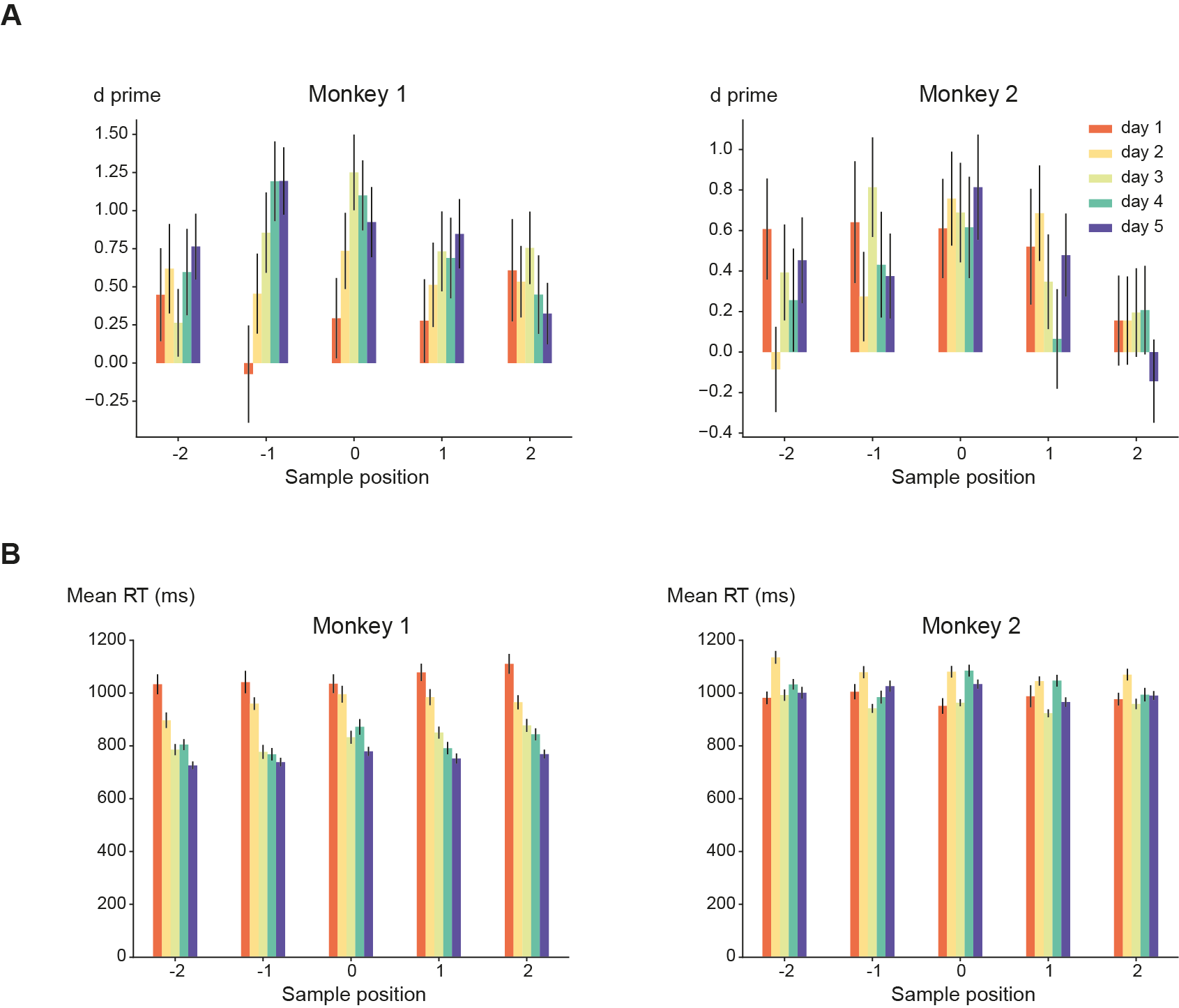

**Figure S2: Behavioural Performance in the match-to-sample task across sample positions**

(A) Discrimination performance (d′ values) for each of the five days of training across different sample positions for monkey 1 (left) and monkey 2 (right). The performance was best when the sample was presented at the fixation point. Error bars indicate standard deviation across 100 bootstrap estimates.
(B) Mean response time (same format as A). Error bars indicate standard deviation **across 100 bootstrap estimates**. Monkey 1’s performance consistently improved over 5 days of training.

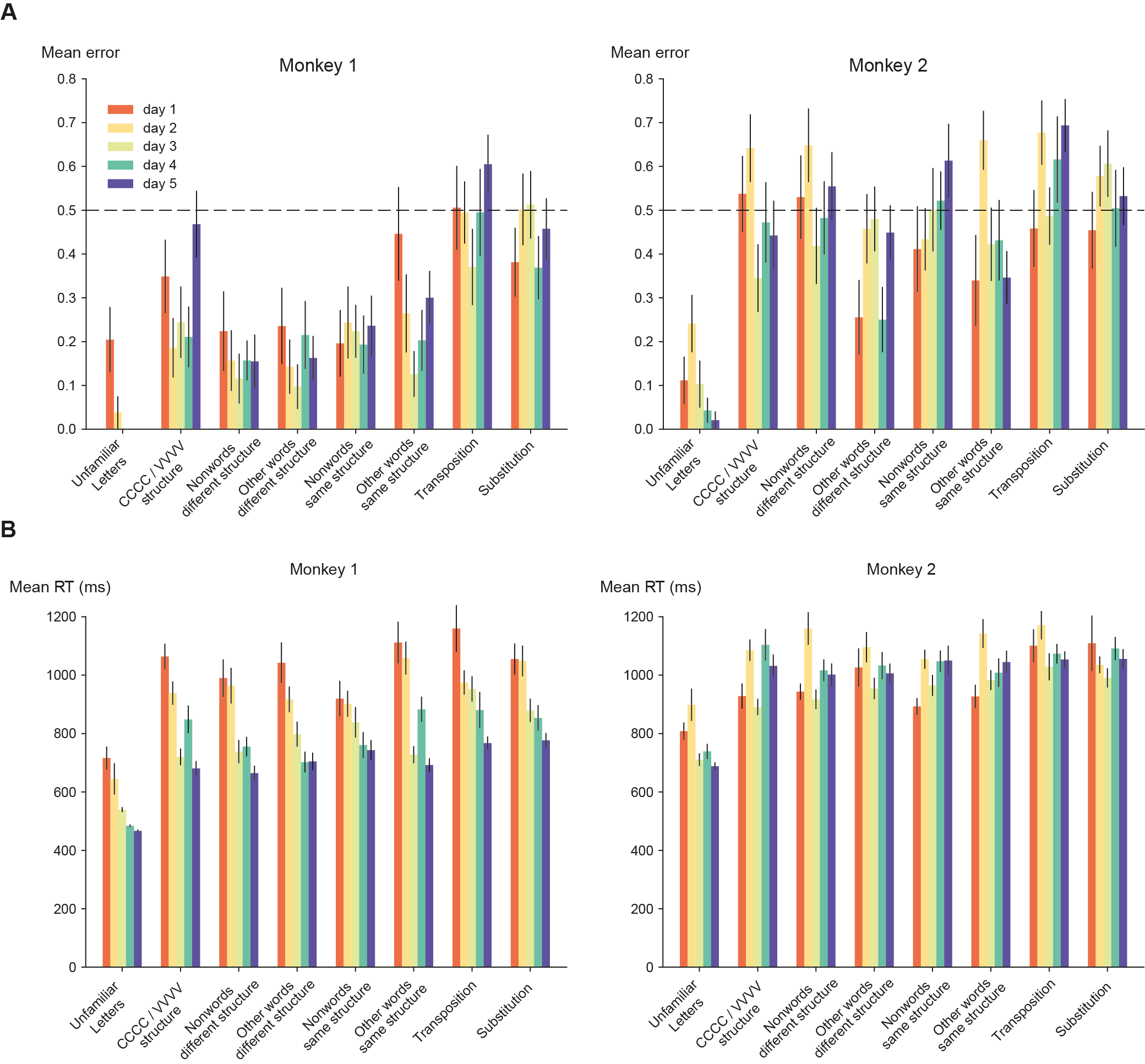

**Figure S3: Behavioural Performance in the match-to-sample task across different nonword conditions**

(A) Mean error rate for each of the five days of training across different nonword conditions for monkey 1 (left) and monkey 2 (right). **Dashed horizontal line indicates chance level. Error bars indicate standard deviation across 100 bootstrap estimates. Both monkeys achieved their best** performance when a different set of letters, never used for writing words, was used (unfamiliar letters, similar to False Fonts in human experiments).
(B) Same as (A) but for mean reaction time. Error bars indicate standard deviation **across 100 bootstrap estimates**. Monkey 1’s performance consistently improved over 5 days of training.

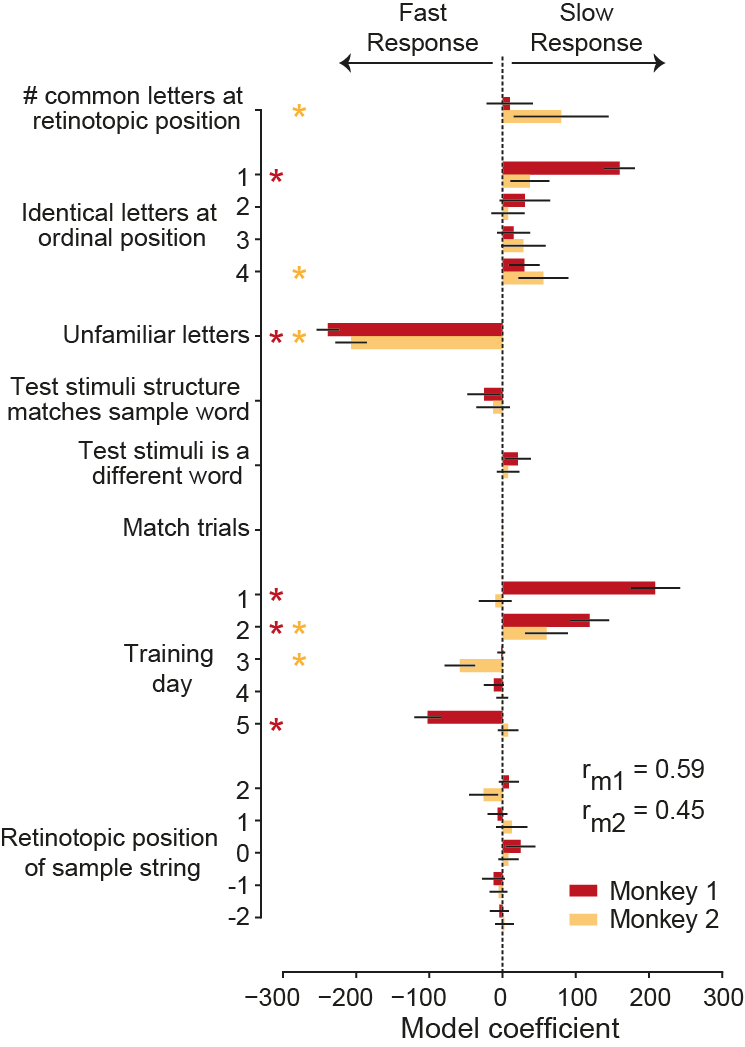

**Figure S4: Reaction Time model fits.** Cross validated (n = 5) LASSO regression model coefficients illustrating factors that lead to "faster" or "slower" response time for monkey 1 (red) and monkey 2 (orange). Error bars indicate one standard deviation across 100 bootstrap estimates. Asterisks indicate statistical significance after Benjamini-Hochberg FDR correction (alpha < 0.05). P-values were estimated from the bootstrap distribution (n = 1000).

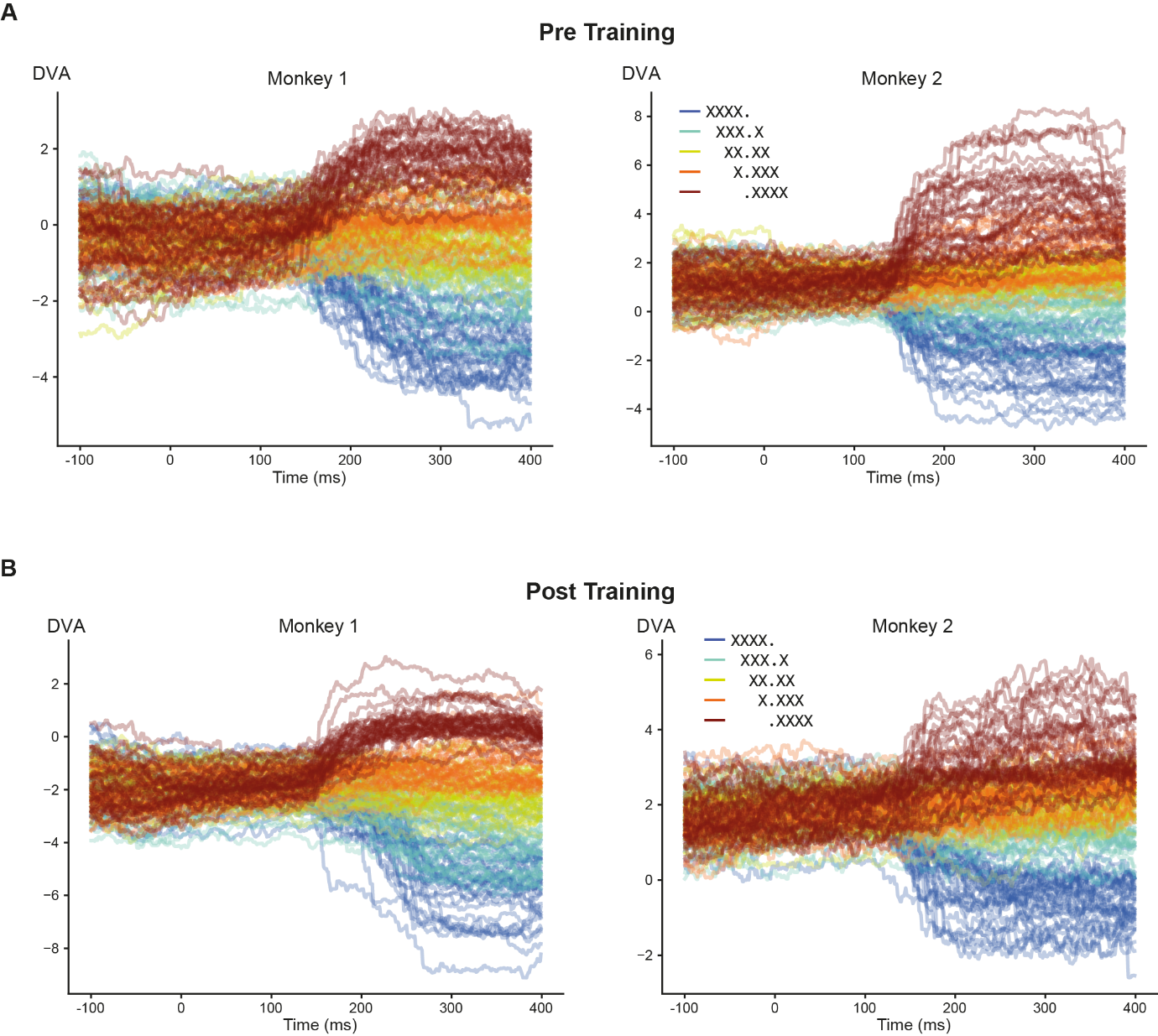

**Figure S5: Eye tracking data during the fixation task**

(A) Mean eye movement traces for Monkey 1 (left panel) and Monkey 2 (right panel). Each line represents the average eye position in degrees of visual angle (dva) over time for a specific stimulus, averaged across all repetitions, and is color-coded according to the sample position of the stimulus. Although stimuli were presented briefly for 200 ms, the monkeys began initiating minor saccadic eye movements approximately 150 ms after stimulus onset, especially for stimuli presented at the most peripheral positions. Note that each letter subtended 4 degrees of visual angle (dva), and the observed eye movements did not exceed the approximate width of one letter, indicating that fixations remained relatively close to the initial point. Furthermore, the stimuli were gone by the time the eye reached its final position (~200 ms).
(B) Same as (A) but for post-training.

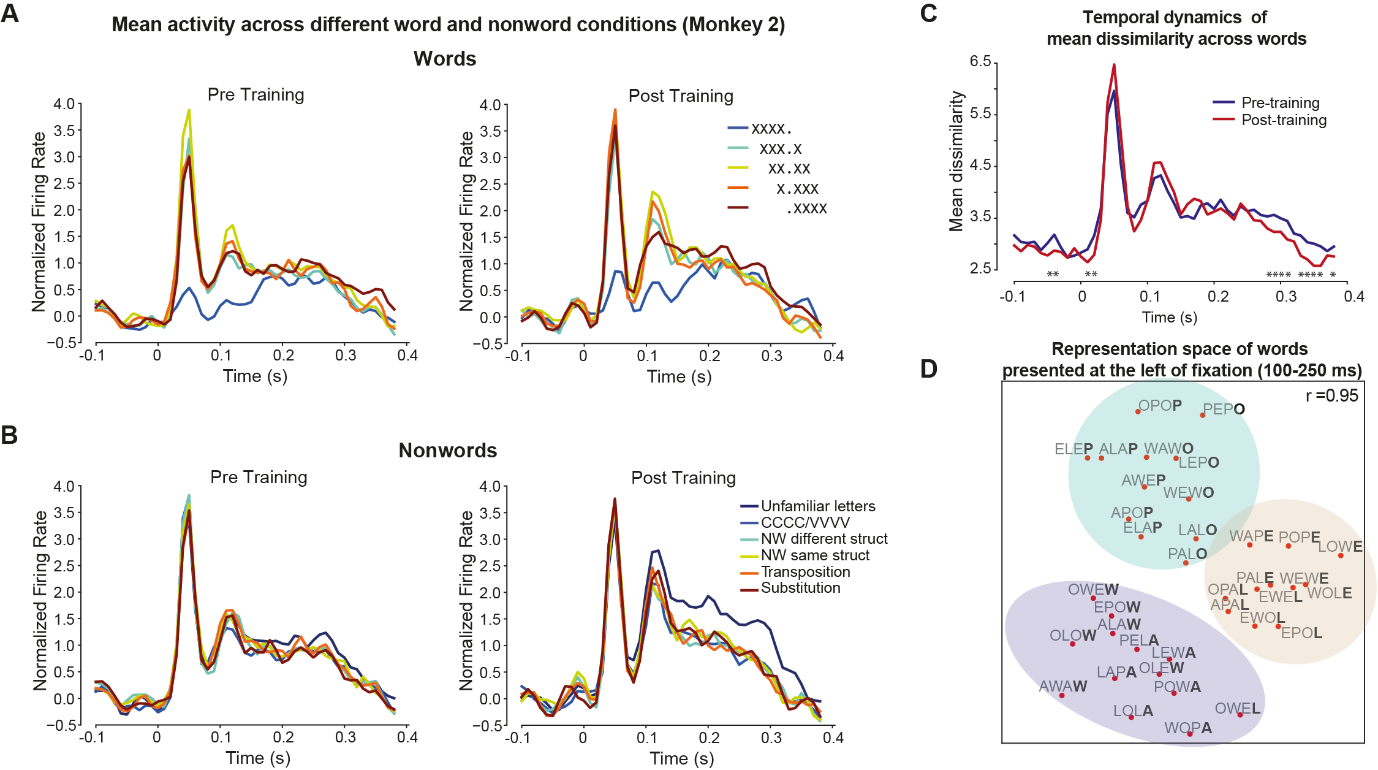

**Figure S6: Average neural population responses and representational similarity analysis (Monkey 2)**

(A) Average normalized neural response over time for words presented across different spatial position during pre (left and post (right) training. The neural responses were highest for the stimuli presented as the center, and minimal for stimuli that fell fully in the contralateral (right) hemifield.
(B) Same as (A) but for different categories of nonwords. We observed a sustained increase in activity for unfamiliar letters post training compared to the other conditions.
(C) Representational dissimilarity estimates pre-training and post-training showing increased dissimilarity among word stimuli post-training. This indicates an expanded neural representational space. Due to a very high degree of freedom (^36^C_2_) all differences are statistically significant.
(D) Multidimensional scaling (MDS) plot of the neural representational space for words (pretraining 100-250ms) presented at the extreme left position, illustrating clustering based on the last letter and letter shape – letters “E” and “L” are clustered together; similarly, for “A” & “W” and “O” & “P”. The last letter of each word is highlighted to emphasize this clustering.

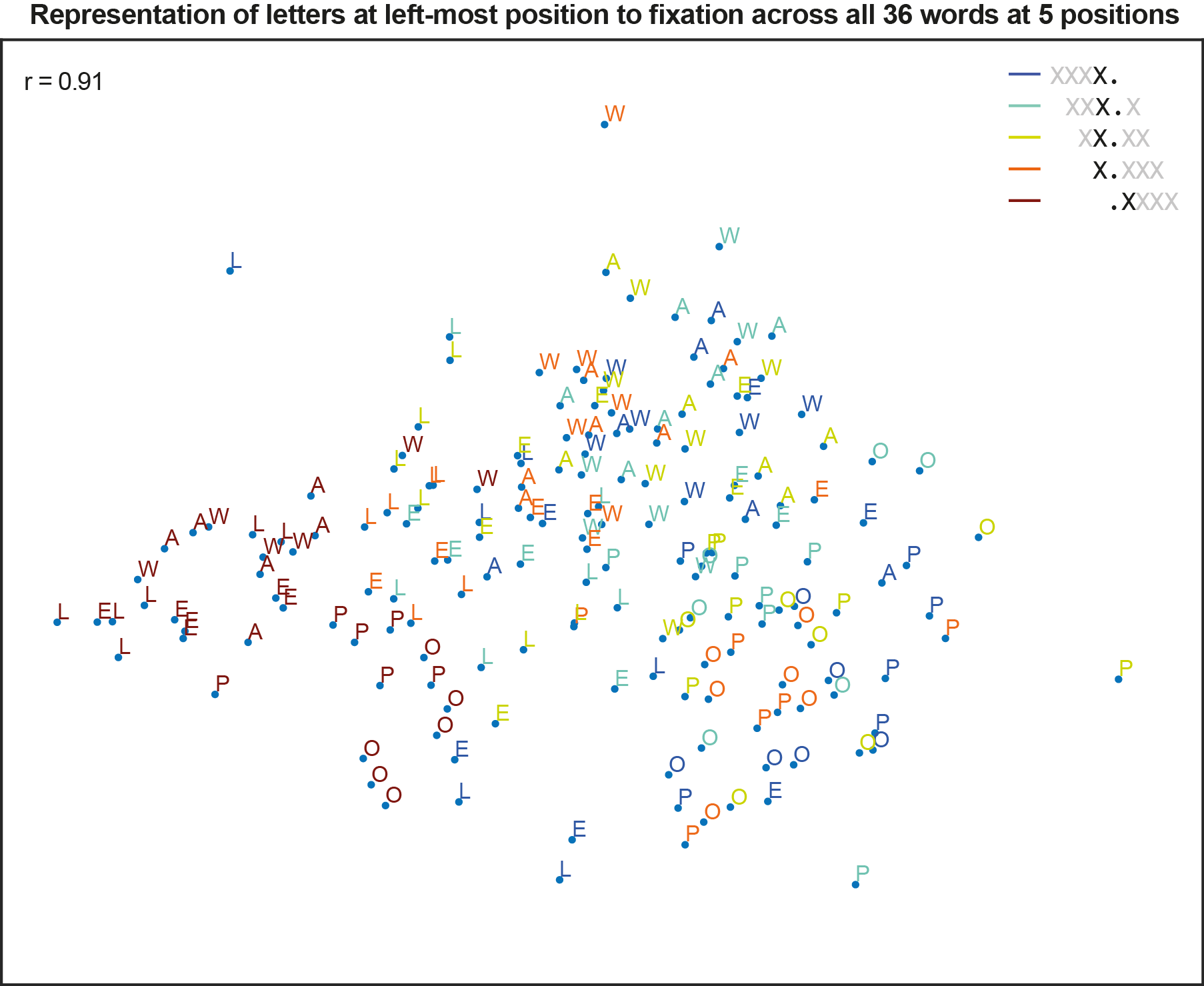

**Figure S7:** **Visualization of the representational space across all word stimuli (Monkey 1)**. The plot shows a multidimensional scaling (MDS) projection of the neural population responses, where each axis represents one of the two principal dimensions capturing the greatest pairwise dissimilarities between neural response patterns. The quality of embedding is estimated by correlating (r) the dissimilarity within this embedded 2d space with the observed dissimilarities from the neural data. For clarity, each data point is labelled with the letter immediately to the left of the fixation point, except for stimuli presented at the extreme right position, where the first letter is shown. The letters are color-coded based on the spatial position of the word. Overall, we observe clustering based on letter identity irrespective of word position. These letter clusters also group according to letter similarity—for example, letters **O** and **P** are positioned closer to each other. However, letters from the extreme left position are separated from the rest of the clusters, possibly due to lower neural responses at that spatial location.

*Representation space of words*

Single-unit analysis revealed that most neurons have retinotopic receptive fields, while ordinal coding explains only a small fraction of the overall variance. To explore these properties at the population level, we modelled the representation space of words as a linear combination of representational dissimilarity matrices (RDMs) corresponding to word position, retinotopic letter position, and ordinal letter position. Specifically, the model RDMs were built by computing the pairwise dissimilarities between all word stimuli using Euclidean distance for each factor derived from the design matrix that was used to predict the firing rates of single units. Thus, the design matrix of the dissimilarity model included 5 factors representing word positions, 8 factors representing retinotopic letter positions, and 4 factors representing ordinal letter positions (Figure S8). The neural data RDM was estimated for each 20 ms time bin across units that showed significant split-half correlations, separately for pre-training and post-training data.

In this linear model Y = Xb; Y is a vector of size (^180^C_2_ x 1) (i.e., pairwise comparisons between 180 stimuli), which indicates the firing rate dissimilarities between pairs of stimuli; X is a matrix of size (^180^C_2_ x 17) excluding the constant term; and b is an unknown weight vector of size (17 x 1). Model coefficients were estimated using 5-fold cross-validated LASSO regularized linear regression (LassoCV function in python). To assess the unique contribution of each factor, we performed variance partitioning like in the single-unit analysis. First, we fitted a model using only the word position predictor and calculated the squared Pearson correlation coefficient (r²) to quantify the variance explained by word position alone. Then, we included the retinotopic position predictors along with word position, recalculated r², and attributed the additional variance explained to retinotopic encoding. Finally, the variance uniquely explained by ordinal position encoding was determined by subtracting the r² of the combined word and retinotopic position model from the r^2^ of the full model.

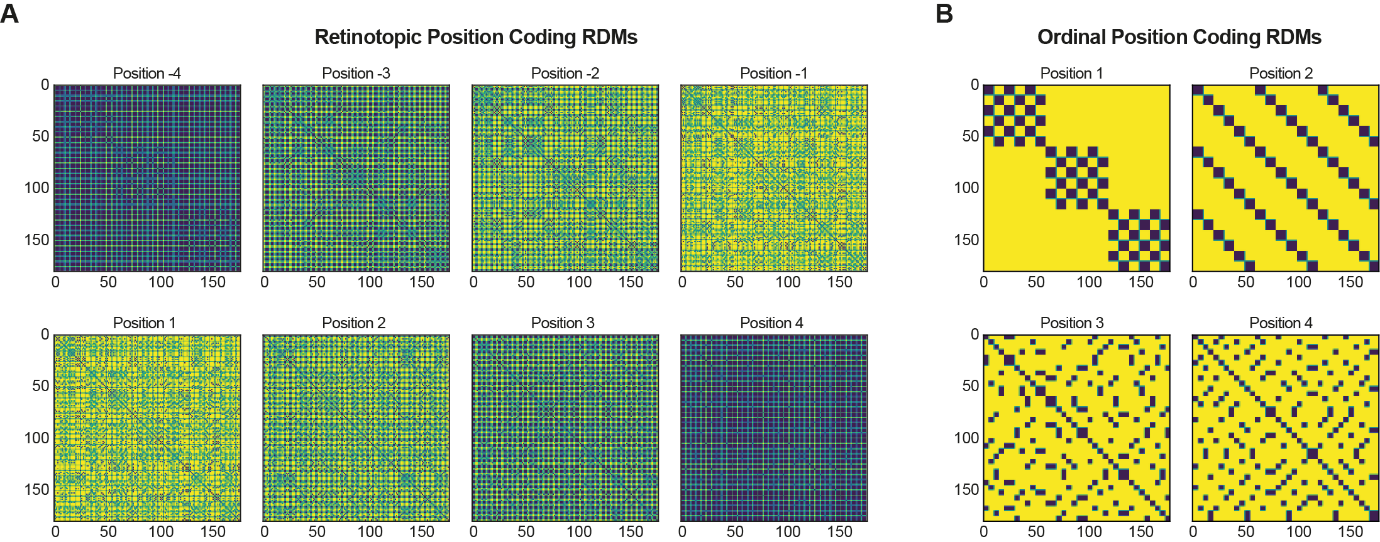

**Figure S8: Model Representation Dissimilarity Matrices (RDM)**

(A) Representational dissimilarity matrices calculated using Euclidean distance for each of the eight retinotopic positions across all 180 word stimuli. Higher dissimilarity values are indicated in yellow, while lower values are shown in blue. Due to the fewer stimuli presented at positions -4 and 4, most of the dissimilarity values at these positions are zero, resulting in less variation within the matrices. In contrast, positions -1 and 1 exhibit a wider range of dissimilarity values, reflecting greater variability in neural representations at these positions.
(B) Same as (A) but for ordinal letter positions.

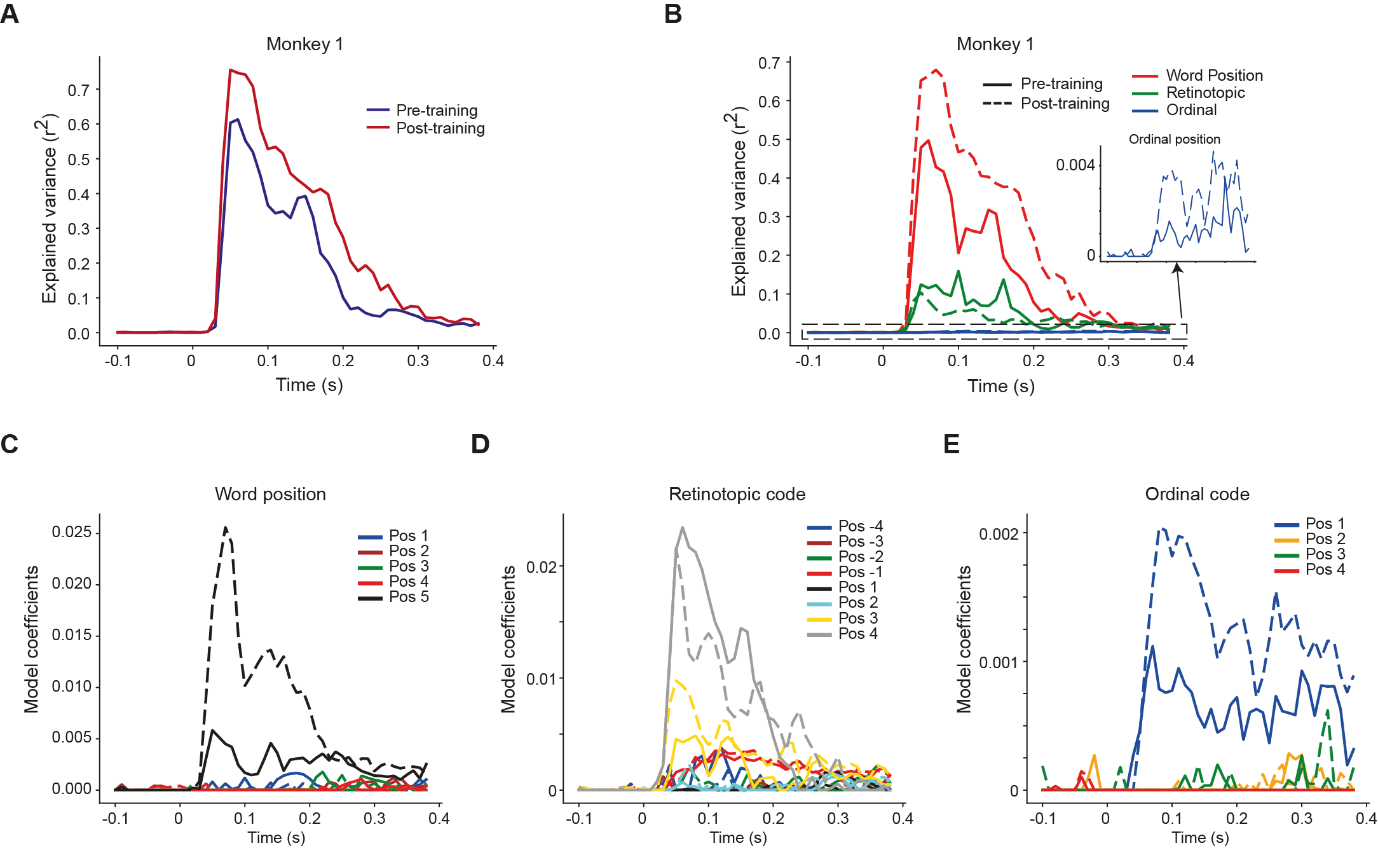

**Figure S9: Dissimilarity model fits for monkey 1**

(A) Proportion of variance explained by the full dissimilarity model at different time points during pre-training (blue) and post-training (red). The better model fits post-training could possibly be explained due to an increase in dissimilarity values.
(B) Proportion of variance explained by the individual components of the dissimilarity model related to word position (red), retinotopic position (green), and ordinal position (blue). The inset magnifies the small but significant variance explained by the ordinal position component. Solid lines indicate pre-training and dashed lines indicate post training. Albeit small, the variance explained by the ordinal position increases after training, indicating enhanced sensitivity to letter order.
(C) Estimated model coefficients for each of the sub-components of the full model, showing their contribution to the overall fit. As in (B), solid lines indicate pre-training and the dashed lines indicate post-training. We observe that word position 5 exhibit the highest coefficient magnitude possibly due to lower activity values at that position leading to higher dissimilarity values. The retinotopic letter positions in the contralateral visual field, especially those adjacent to the fixation, emerged as a key predictor of neural representation space, along with the first ordinal position.

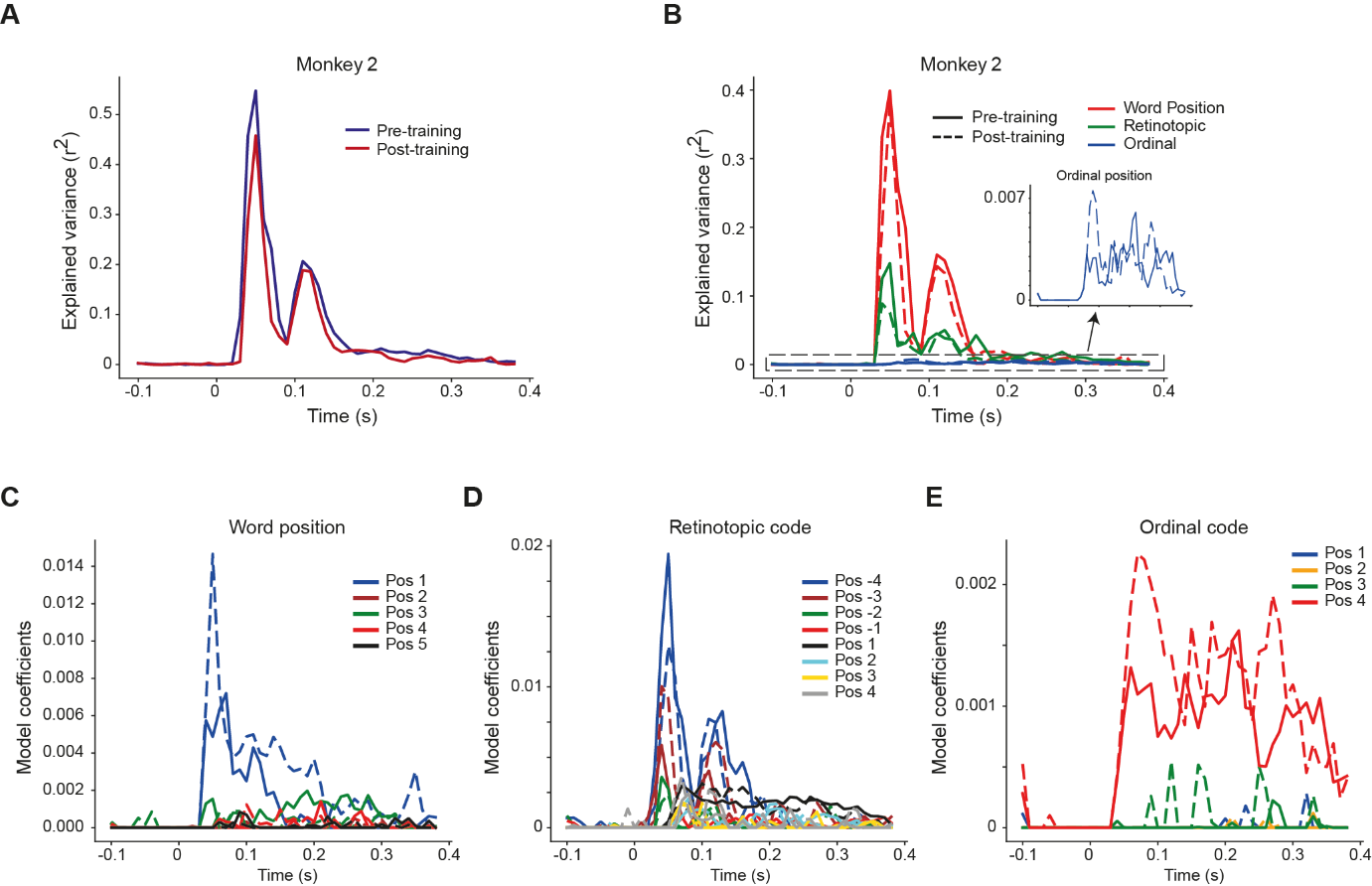

**Figure S10: Dissimilarity model fits for monkey 2**

(D) Proportion of variance explained by the full dissimilarity model at different time points during pre-training (blue) and post-training (red). The change in model fit across time parallel the changes in mean neural activity (see figure S6).
(E) Proportion of variance explained by the individual components of the dissimilarity model related to word position (red), retinotopic position (green), and ordinal position (blue). The inset magnifies the small but significant variance explained by the ordinal position component. Solid lines indicate pre-training and dashed lines indicate post training.
(F) Estimated model coefficients for each of the sub-components of the full model, showing their contribution to the overall fit. As in (B), solid lines indicate pre-training and the dashed lines indicate post-training. Similar to Figure S7, we observe that word position 1 exhibit the highest coefficient magnitude possibly due to lower activity values at that position, leading to higher dissimilarity values. The retinotopic letter positions in the contralateral visual field, especially those adjacent to the fixation, emerged as a key predictor of neural representation space, along with the last ordinal position. Notably, word position was dominant only in the initial time window (0-200 ms)

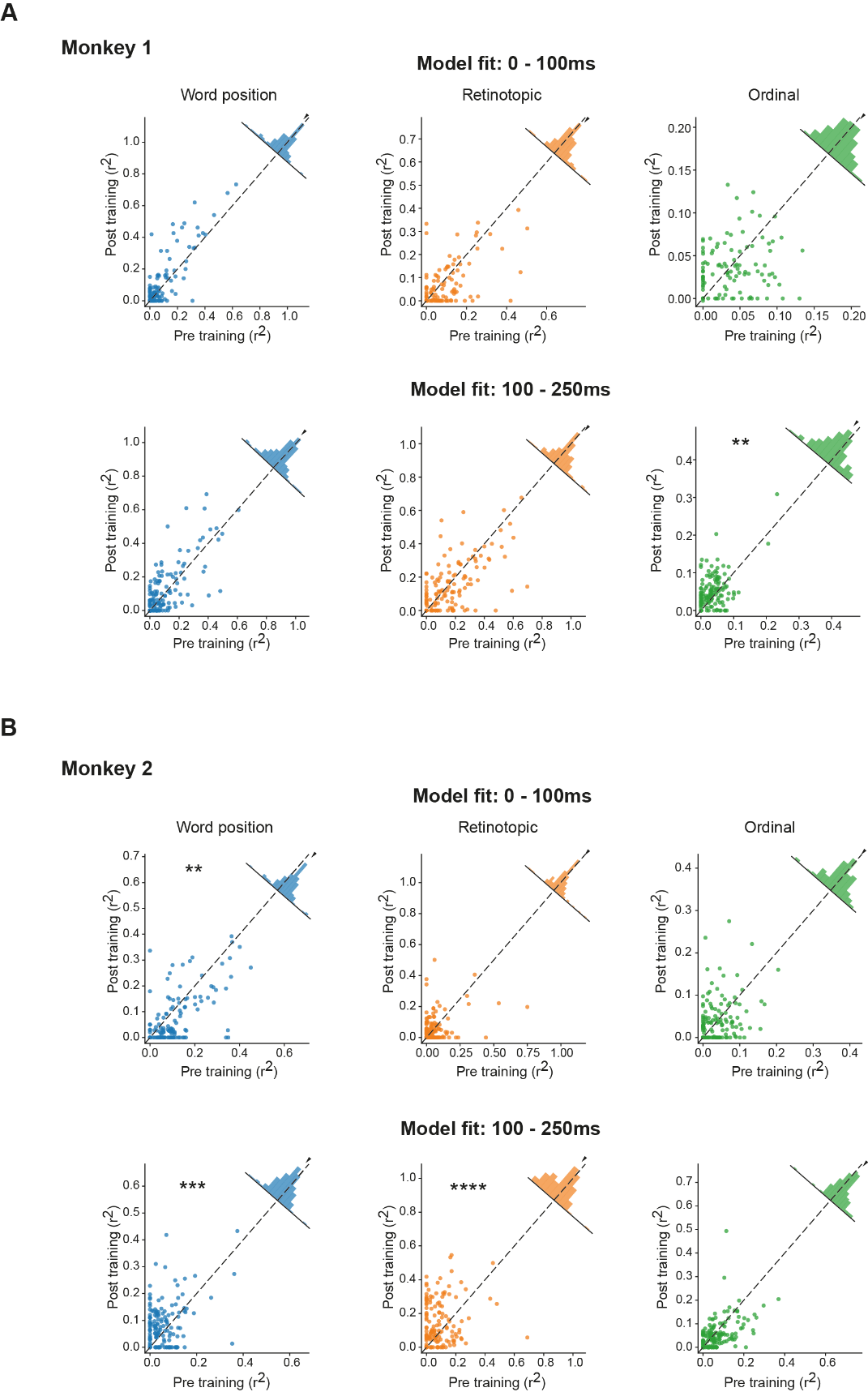

**Figure S11: Distribution of model fit performance across all neurons**

(A) Proportion of variance explained by word position, retinotopic, and ordinal factors during early (0–100 ms) and late (100–250 ms) time windows for monkey 1. Each dot represents a single neuron that was recorded both before and after training. Dashed line represents unity slope. The difference (post-pre) across all neurons is shown using histogram along the unity slope line and the mean value is indicated via triangle. Asterisks indicate statistically significant increase in post training unless otherwise stated (**, p< 0.005; ***, p<0.0005, and so on, paired t-test).
(B) Same as (A) but for monkey 2.

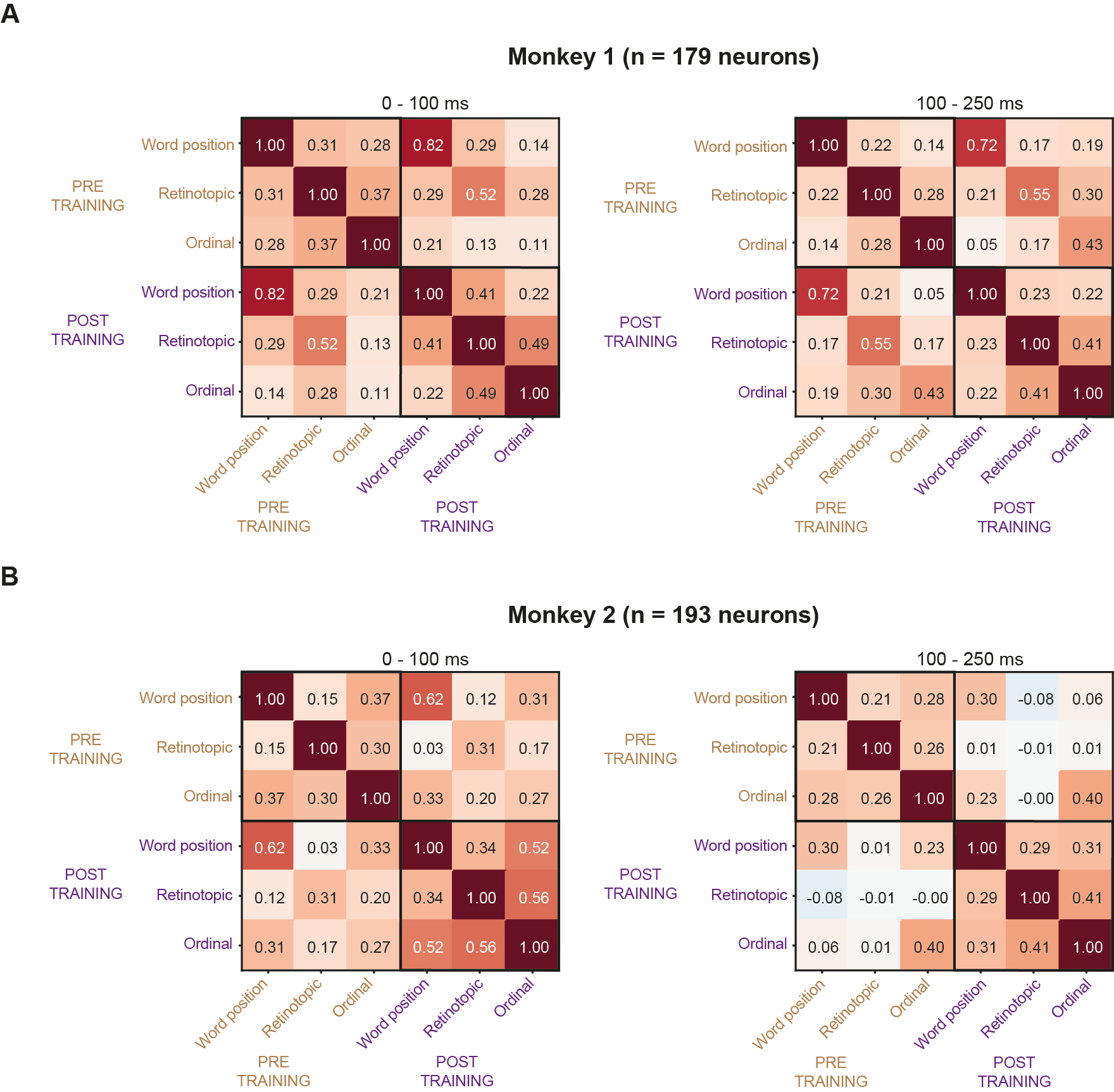

**Figure S12: Correlation of feature encoding strength before and after training.**

(A) Pair wise correlation between the variance explained by each of the independent components of the letter x position model before and after training for monkey 1. The correlation matrix is computed for the early (left) and late (right) time windows. High off-diagonal correlation values suggest that the tuning properties that individual neurons possessed before training remained largely preserved post training, even though a larger number of neurons encoded ordinal letter position after training (Figure S11A).
(B) Same as (A) but for monkey 2.

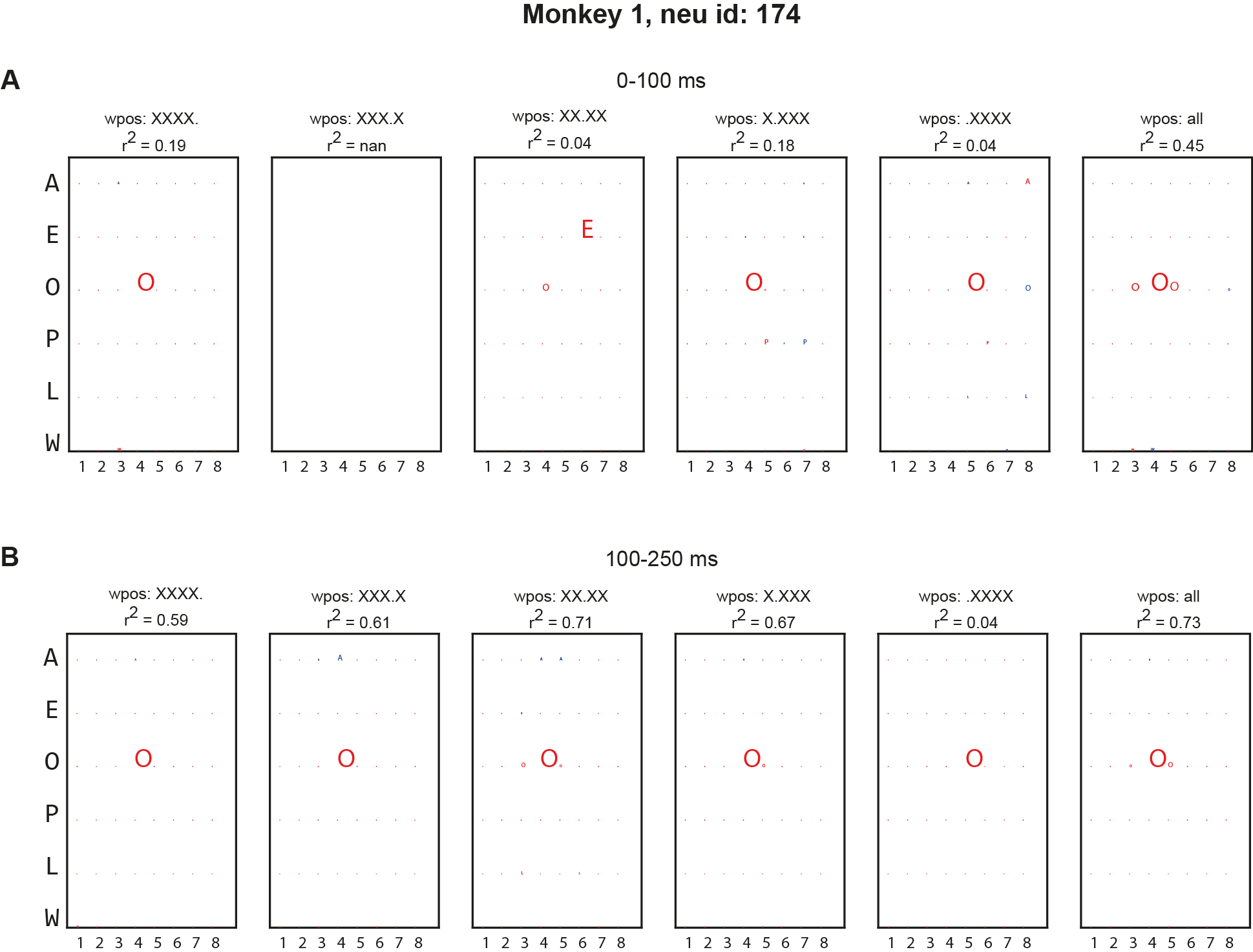

**Figure S13: Estimated retinotopic receptive field of unit 174 (pre-training)**

(A) Model coefficients estimated from the LASSO regression trained independently on stimuli presented at each word position (wpos) during the early (0-100ms) time window. The x-axis indicates the possible retinotopic position of each letter within the word, and the y-axis lists the individual letters. Each panel shows the estimated receptive field weights, reflecting how strongly neural responses are modulated by specific letter identities at given spatial locations. The overall variance explained (r^2^) for each model is shown above each panel along with corresponding word position (wpos). The estimated receptive field remain fairly consistent across word positions. In the ipsilateral visual field, the model fits are weaker, likely due to lower activity, and the estimated receptive field is shifted towards the fixation point while preserving the preference for letter identity. The final (rightmost) panel shows the overall model fit when data from all five word positions are combined.
(B) Same as (A) but for the late time window (100-250ms). We observe no change in the receptive field due to subtle eye movements when stimuli were presented across different spatial positions.

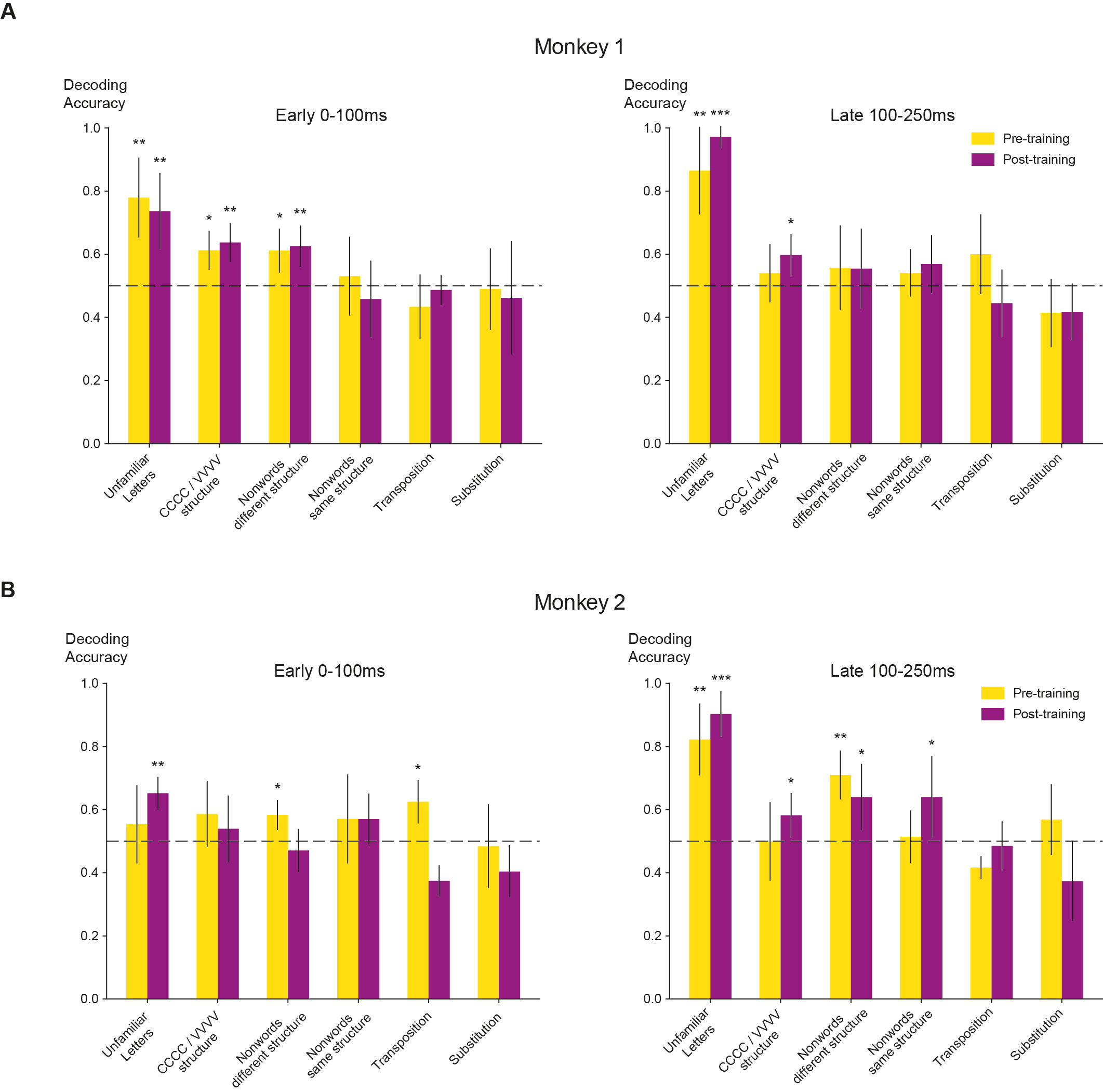

**Figure S14: Neural decoding of word vs nonwords**

(A) Cross-validated decoding accuracy in distinguishing words from each of different categories of nonwords (x axis), separately for early (left) and late (right) time windows in Monkey 1. Error bars represent the standard deviation across five folds. Asterisks denote significance levels (*p < 0.05; **p < 0.005; ***p < 0.0005; one-sample t-test relative to the chance level of 0.5). Decoding accuracy for unfamiliar letters increases during the late time window, whereas decoding based on structural differences is primarily confined to the early time window.
(B) Same as in (A), but for Monkey 2. Prior to training, mean neural responses to unfamiliar letters were comparable across nonword types; nonetheless, words and nonwords comprised of unfamiliar letters could still be discriminated during the late time window. Following training, this distinction emerged earlier, becoming detectable in the initial time window as well.

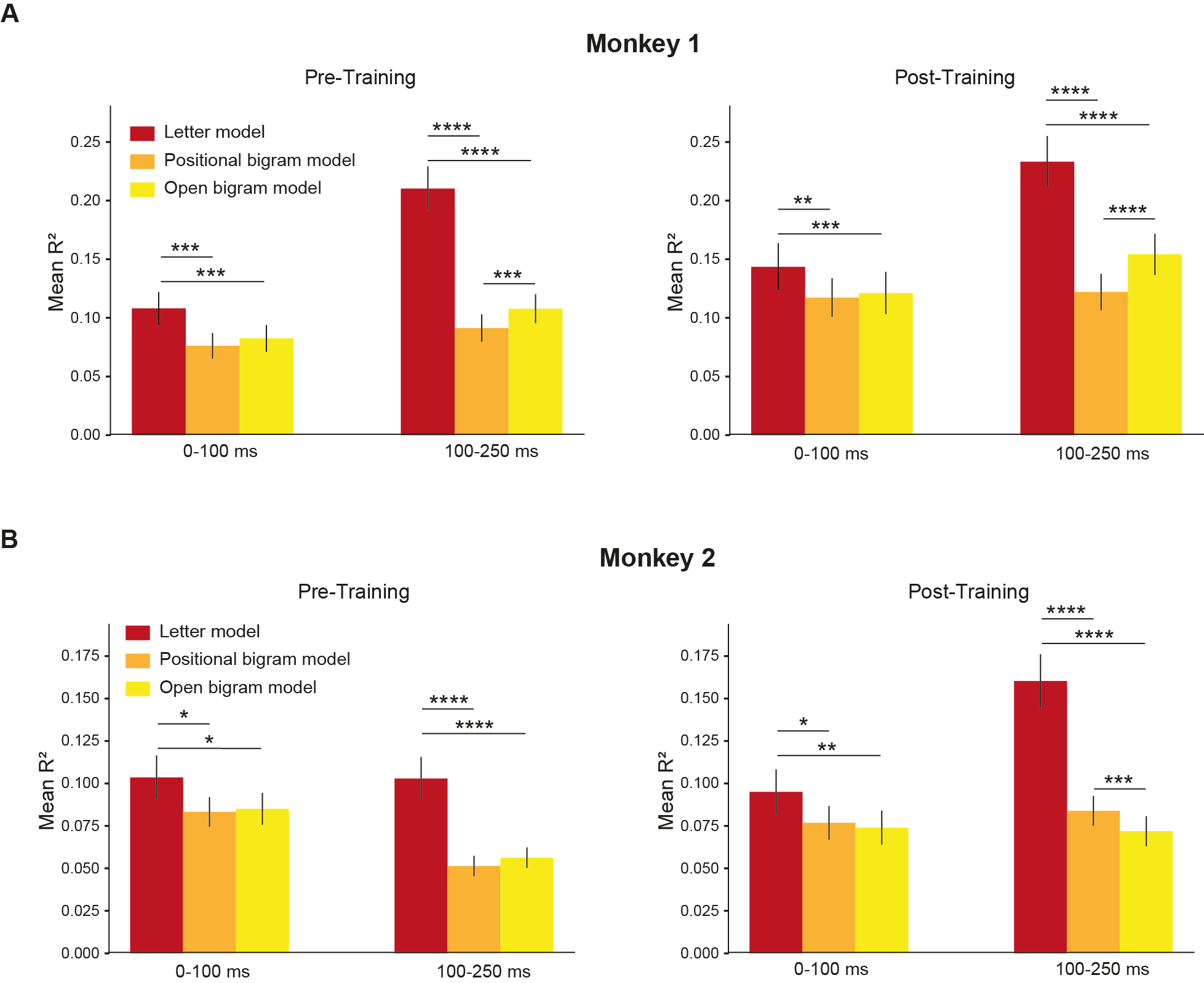

**Figure S15: Comparing letter vs bigram model**

(A) Mean explained variance (R^2^) across all IT neurons in Monkey 1 with significant split-half reliability, shown for three models: letter x position model, position constrained bigram model (using CV or VC bigram only), and the open bigram models. The letter x position model fits improved during the late time window (100-250ms) compared to early time window (0-100ms) and significantly outperformed both bigram models in pre-training (left) and post-training (right) sessions. Error bars indicate standard error of mean. Asterisks indicate statistical significance (*, p< 0.05; **, p< 0.005, ***, p<0.0005, and so on, Paired t-test)
(B) Same as (A) but for monkey 2
